# Evaluation of a structure-based method for *ab initio* gene detection using deep learning

**DOI:** 10.64898/2025.12.19.694709

**Authors:** Jonathan Hummel, Karol Estrada

## Abstract

In this work, a novel method for the detection of exons within genomic DNA sequences was implemented and evaluated. This is a structure-based approach, inspired by recent work from Sharma et al., by which nucleotide sequences are converted into physicochemical profiles based on trinucleotide and tetranucleotide mappings that were estimated via molecular dynamics simulations [8,12]. Three deep learning models with architectures suitable for multidimensional sequence classification were trained on the structural profiles for sequences at the junctions between exons and introns, as well as those sampled from other regions of a human reference genome. The trained models showed promising performance in the evaluation set, but will require a much more sophisticated and robust post-processing approach to achieve compelling exon- or gene-level classification performance in realistic use cases. A tool by Sharma et al. based on the same structural approach, which they named ChemEXIN [12], was also evaluated. The results indicate that the approach taken in this work during the development and application of the models offers a major improvement over ChemEXIN, and may more accurately reflect the potential of the underlying idea.

## 1) Introduction

The computational detection of genes, protein-coding or otherwise, within eukaryotic DNA sequences has been a challenging problem for decades. Lower gene density and introns/splicing make it particularly difficult to identify a gene’s structure accurately in eukaryotes, compared to prokaryotes [1]. Early solutions to this challenge relied on homology with known genes and sequences. While this approach is well-grounded and certainly has its strengths, when used alone, it can fail if the target sequence does not have sufficient significant matches among the reference genome(s). Also, accurately inferring gene structure from homology hits alone presents further challenges [1]. *Ab initio* gene detection methods aim to use more general and abstract features of a gene’s typical structure to identify the likely location of genes within a sequence. These types of methods have traditionally relied on encoding and using information about key locations, such as splice sites, branch points, and start codons, as well as species-specific patterns found in the coding regions themselves [1].

The first wave of gene detection programs only aimed to predict the locations of exons, with no further context. After that came a wave of programs that attempted to predict a gene’s full structure. These were algorithmically diverse solutions, including GeneID, GeneParser, and FGENESH, among others [1]. These early programs relied on the strong assumption of one complete gene per input sequence, and they typically exhibited poor performance. First GENSCAN, and then AUGUSTUS, each offered major improvements in performance compared to previous solutions and were both based on versions of the generalized hidden Markov model (HMM) [1,2]. AUGUSTUS probabilistically models the sequences that make up critical parts of a protein-coding gene’s structure, including the splice site, the branch point, initial coding bases, coding regions, and non-coding regions, among others. Part of the strength of AUGUSTUS comes from its detailed modeling of exon, intron, and intergenic length distributions, which are complex and positionally dependent [2]. While HMMs have traditionally been one of the most successful approaches to this problem, they often encode information about gene structure that may not be accurate or available for sequences from newly sequenced genomes [1]. That said, AUGUSTUS has been widely used and well regarded for the past two decades. BRAKER is a gene annotation pipeline that uses AUGUSTUS, along with several other tools and reference sequence sources. BRAKER3 is currently state-of-the-art for accurate gene detection in new eukaryotic genomes and is a hybrid of *ab initio* and sequence homology-based methods [3,4].

Some of the earliest gene detection programs were based on neural networks, but these were not very competitive [1], likely because the theory and compute power at the time were insufficient for such a challenging problem. However, deep learning has recently been applied to gene detection quite successfully, and examples include Helixer, ANNEVO, and Tiberius [4–6]. Helixer combines convolutional neural network (CNN) layers with bidirectional long short-term memory (BiLSTM) layers to, once trained, classify each base as either part of a coding sequence (CDS), intron, untranslated region (UTR), or intergenic region. An HMM converts these nucleotide-level predictions into the most probable gene structure that adheres to known biological rules (via the Viterbi algorithm). Helixer is a purely *ab initio* method, and it achieved good generalizability across diverse species and competitive performance compared to AUGUSTUS [6]. Like Helixer, Tiberius also combines CNN and BiLSTM layers to learn abstract representations of genetic features from sequences, along with an HMM to find the most probable gene structure from among the deep learning layers’ predictions. Unlike Helixer, the HMM model in Tiberius is integrated and trained with the deep learning layers. It uses a more efficient implementation of the Viterbi algorithm, and calls more classes, allowing for a more nuanced consideration of gene structure. Tiberius was also trained with a special loss function to address the class imbalance in gene detection [4]. The authors of Tiberius built upon and notably improved the already impressive precedent set by Helixer. As such, it is unsurprising that it performed competitively when compared to existing state-of-the-art tools, including AUGUSTUS and BRAKER3, and that it generalized well across animal species, especially at the exon level [4,6]. ANEVO, which is still under review, uses a different deep learning approach called mixture-of-experts. In this approach, a collection of deep learning networks is trained, with each model intended to detect specific qualities of a gene’s structure. The output of each model is dynamically weighed/gated using genomic context to arrive at the final predictions. ANEVO was trained on the genomes of many diverse organisms simultaneously to learn evolutionary relationships, which enables excellent generalizability across species. ANEVO performed competitively when compared to AUGUSTUS, BRAKER3, and Helixer [5].

In 2021, Mishra et al. published their findings that, on average, unique deviations in the B-DNA structure occur around splice sites in the human genome. This pattern was observed in physicochemical parameters that describe the structure of the DNA helix, which were estimated from the dimers in genomic sequences using data extracted from X-ray crystallography structures [7]. This observation from Mishra et al. was verified in the early stages of the work detailed here (see ***Figure 6***). The meaning of some of these structural parameters can be found in ***Figure 1***. In 2023, the same research group published a paper showing that similar deviations in the DNA structure exist on average, not only around splice sites in humans but also around the splice sites and transcription start sites (TSS) in 8 yeast species. They observed a similar pattern around the TSSs of 12 prokaryotic species as well [8]. In that paper, they used a more advanced method, involving molecular dynamics simulations, to estimate the physicochemical parameters as a function of DNA trinucleotides and tetranucleotides, rather than dinucleotides [8]. These papers were interesting for two reasons. First, as the authors emphasized, the predicted existence of conformational signatures at key locations that define the layout of genes might offer a new modality, beyond the sequence space, by which genes could be reliably identified *ab initio* in genomes across diverse species [7, 8]. Second, these observations may provide a basis for a more detailed understanding of splicing. On one hand, the frequent deviation in the B-DNA structure around exon junctions may be a sort of “complementary shadow” of the structural features in the corresponding pre-mRNA that are more directly relevant to splicing. That said, *in vivo*, splicing largely occurs co-transcriptionally, and there is evidence that the rate of elongation of the nascent transcript may affect the splicing machinery and splicing outcomes [7,9–11]. As such, it is not a major stretch to imagine that conformational properties of the genomic DNA itself could have some type of effect on splicing by modulating the elongation rate of pre-mRNA at specific locations within a gene.

**Figure 1:**
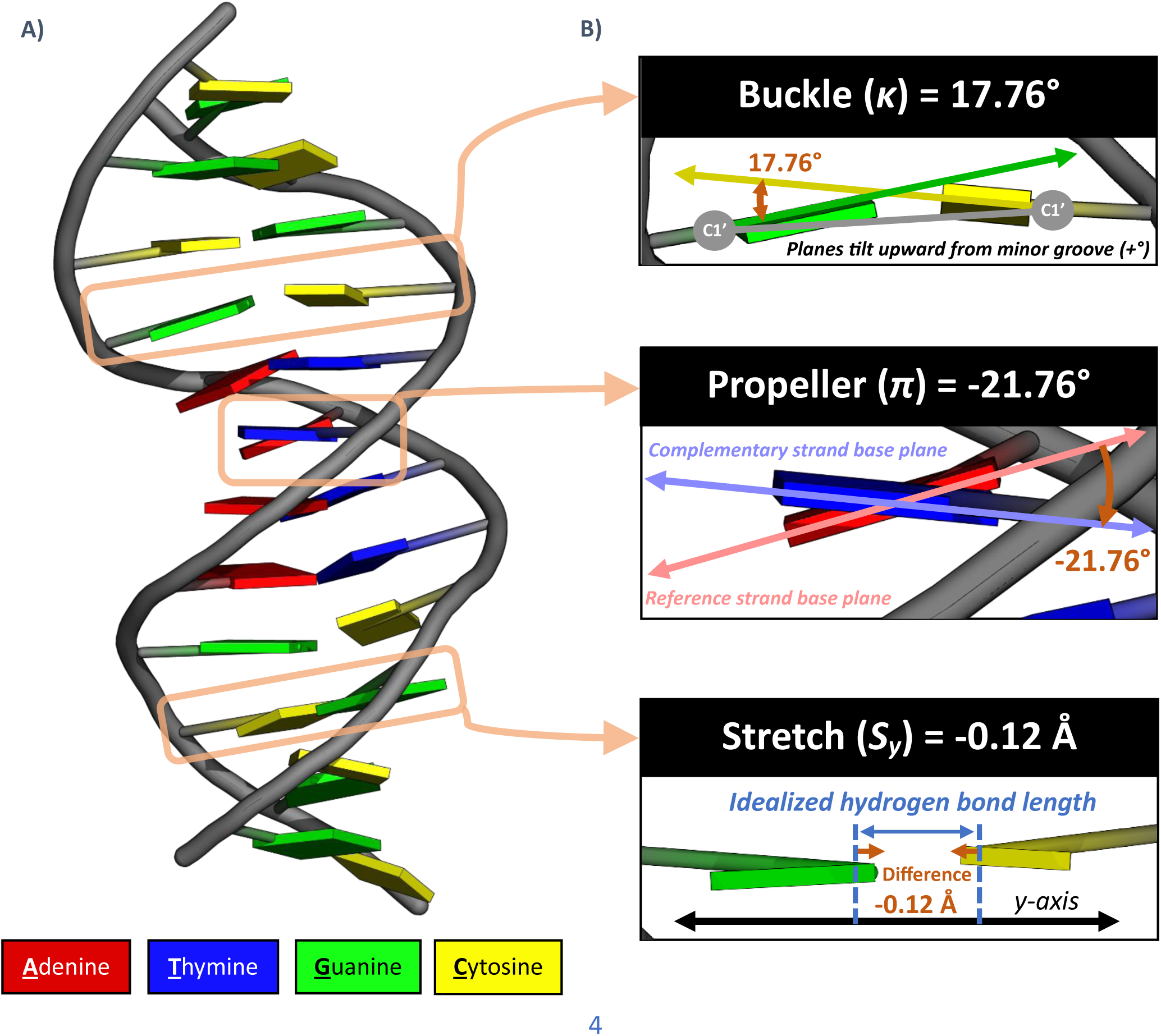
**A)** Cartoon representation of a B-DNA helix, with purine and pyrimidine planes represented as color-coded rectangular prisms. Rendered in PyMOL with the help of X3DNA-DSSR (Protein Data Bank structure 4BNA) [13,14]. **B)** A demonstration of three of the structural parameters used by Sharma et al., as well as in the work detailed below. These are all intra-base pair parameters, but many additional parameters describe the backbone and inter-base pair step characteristics, among other things [12,15]. The parameter values reported below were calculated for this structure using X3DNA-DSSR. The reference lines and markers are exaggerated for clarity, and are only intended to give conceptual context. Please see Figure 6 for an enumeration of all 28 parameters used in this work. Also, see the review by Da Rosa et al. for more depictions and context regarding the parameters and conventions used to describe DNA structure [17].

Unfortunately, investigating the latter idea is outside of the scope of this work, and would best be conducted in a lab. Instead, the work described below aims to explore whether these structural signals around the junctions between introns and exons show any promise for *ab initio* gene detection via deep learning. This year, Sharma et al. published a paper about their effort to build upon their previous work cited above to do just that. That work culminated in a command-line tool, which they named ChemEXIN [12]. The state-of-the-art tools described earlier were all trained on sequence data directly. The CNN model at the heart of ChemEXIN was instead trained on multidimensional structural profiles derived from sequence data and molecular dynamics simulations. These numerical profiles were based on parameters that can be used to describe the structure and properties of B-DNA as a function of sequence. Please see ***Figure 1*** for an example of the types of properties these parameters represent. As such, ChemEXIN has been trained on data in the structural domain. While Sharma et al. referred to these feature sets as “physicochemical profiles” [12], in this work they are defined as “predicted structural profiles” and hereafter referred to simply as “structural profiles”. This distinction is made because the majority of the parameters (25 of 28) correspond to the geometric characteristics of the double helix, while the remaining three are associated with its thermodynamic stability (see ***Figure 6***). Furthermore, the profiles are moving averages of values obtained via trinucleotide and tetranucleotide lookup tables derived from molecular dynamics simulations, rather than direct experimental data. According to the paper by Sharma et al., ChemEXIN performed much better than traditional tools, including AUGUSTUS, GENSCAN, GeneID, and FGENESH [12]. That said, both BRAKER and Helixer were absent from this comparison. A version of BRAKER would have been available and well-known when that research was being conducted, and Helixer likely was as well. When choosing, training, and evaluating the models discussed below, an approach was taken that was inspired by the authors’ original underlying idea, but not identical. As such, this work serves as an independent technical audit of not just the ChemEXIN tool, but also the underlying idea behind it.

## 2) Materials & methods

This work was largely conducted with code written in Python, along with some Bash commands and bioinformatics command-line tools. Specific tools and Python libraries are named and discussed below whenever relevant.

### 2.1) Dataset

The primary data set used in this work was the human genome assembly GRCh38.p14, along with its corresponding RefSeq annotation [16,17]. Both the FNA and GFF files were downloaded from the NCBI website. More specifically, the locations and sequences of exons in protein-coding genes in the training and testing sets were retrieved using the RefSeq MANE Select exon annotation. This subset of annotations was chosen because it includes curated canonical transcripts for every human gene, and these all have exact counterparts in the Ensembl/GENCODE annotation [17]. This helped to avoid repetitive data in the training and testing sets, and enabled easy mapping to Ensembl/GENCODE if ever needed.

Sharma et al. used molecular dynamics simulations to estimate the values of 22 structural parameters of DNA associated with every possible trinucleotide sequence (i.e. trimer). They also did so for every possible tetranucleotide sequence for 6 additional structural parameters. The resulting mappings were shared in the <u>supplemental data</u> for their most recent paper [12]. These tables were used to map trimers and tetramers in the input sequences to their corresponding structural parameter values during the profile generation step, described in **Section 2.3**. Unfortunately, there was not enough time to recreate/validate these mappings independently using available structural bioinformatics tools, as this would have been a major undertaking. See ***Figure 1*** for an illustration of the physical meaning of a few of these parameters within the context of the DNA helix. A review by Da Rosa et al. provides a more in-depth discussion of the structure of DNA and conventions used to describe it [15].

The only other dataset used in this work was a plot of the size distribution of human exons from a paper by Mokry et al. [18]. Data points were taken from Figure 1 of that paper using an online tool that facilitates extracting data points from images of graphs. The resulting data set served as a quick stand-in for more sophisticated model post-processing techniques, which is discussed further in **Section 2.6**.

### 2.2) Sequence sampling

Samtools faidx was used to index the GRCh38 FNA file, and all sequences were retrieved from all chromosomes of GRCh38 using samtools faidx [19], which was called from Python via the subprocess library. As such, the sequence sampling process and organization of data relied on the associated sets of genomic coordinates. All sequences used in training and testing were always tied to their source genomic coordinates, which provided transparency and enabled visual validation of the sequence retrieval and processing scripts using Integrative Genomics Viewer (IGV). All viable sequences for model training, validation, and testing were identified and retrieved from GRCh38 first, and then partitioned later. The sequences retrieved were each 103 bp long. As is described in the next section, the profile generation algorithm calculates values for each position in a sequence using a centered 27 bp sliding window to calculate a moving average for each parameter (see ***Figure 3***). 103 bp gives the algorithm enough sequence to calculate the moving averages for a centered 77 bp subset of the original sequence, because 13 bp are lost on either end of the 103 bp sequence during profile generation. An odd length was chosen because an odd number made it easier to plan and test a center-oriented, window-based algorithm. 77 was chosen based on the average observed behavior around exon start and end sites in the training set, as it provided a safe yet compact window around where the signals likely occur (see ***Figure 6***). This ensured that each model could learn most of the relevant patterns that occur around the exon junctions. 103 bp sequences were sampled from the forward and reverse stands using the RefSeq MANE Select exon annotation via the following criteria: intron-exon (start of a feature), exon-intron (end of a feature), control exon (within a feature), control intron (between subsequent features of the same Gene ID), control intergenic (between subsequent features of different Gene IDs). The sampled 103 bp boundary sequences were allowed to overlap by 13 bp at most, which is the part of the sequence lost to profile generation and which only has a partial influence on the signal for the first or last 13 bp of the 77 bp training window. This meant that boundaries were not sampled from exons shorter than 90 bp. A more stringent padding was enforced between boundary and control sequences, as an exon had to be at least 216 bp long for a sequence to be sampled from its center. This meant that a 103 bp control sequence sampled from the smallest allowed exon would still have ∼5 bp of completely non-overlapping padding between the two adjacent boundary sequences (pre-profile generation). Multiple control sequences were sampled from a single exonic, intronic, or intergenic region by taking as many evenly separated 103 bp sequences as the region would allow, up to a maximum, while adhering to the rules for separation from boundaries detailed above, as well as separation from each other (> 5 bp). The maximum number of control sequences that could be sampled from a single region was 60 from an exon, 20 from an intron, and 180 from an intergenic region.

**Figure 2:**
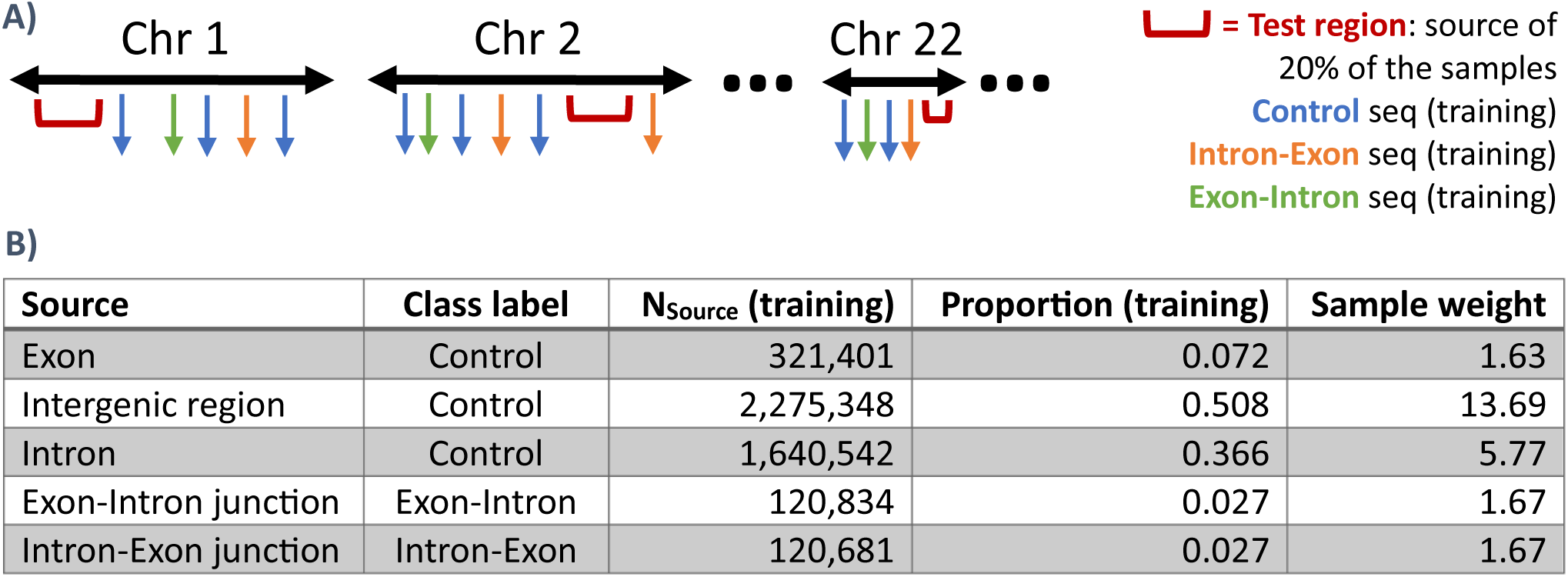
**A)** Overview of sampling strategy to ensure contiguous regions of each chromosome remained unseen to the models during training. This enabled longer test sequences to be drawn after testing for more rigorous evaluation. **B)** The final total counts of each type of profile in the training set after profile generation (see next section). Each profile corresponded to a 77 bp sequence. Proportions of each sequence source are shown for the training set, but were approximately equal in the training and testing sets (sum squared error of 0.002). Sample weights were used during training to prepare the model for use within the context of a genome (see explanation below).

**Figure 3:**
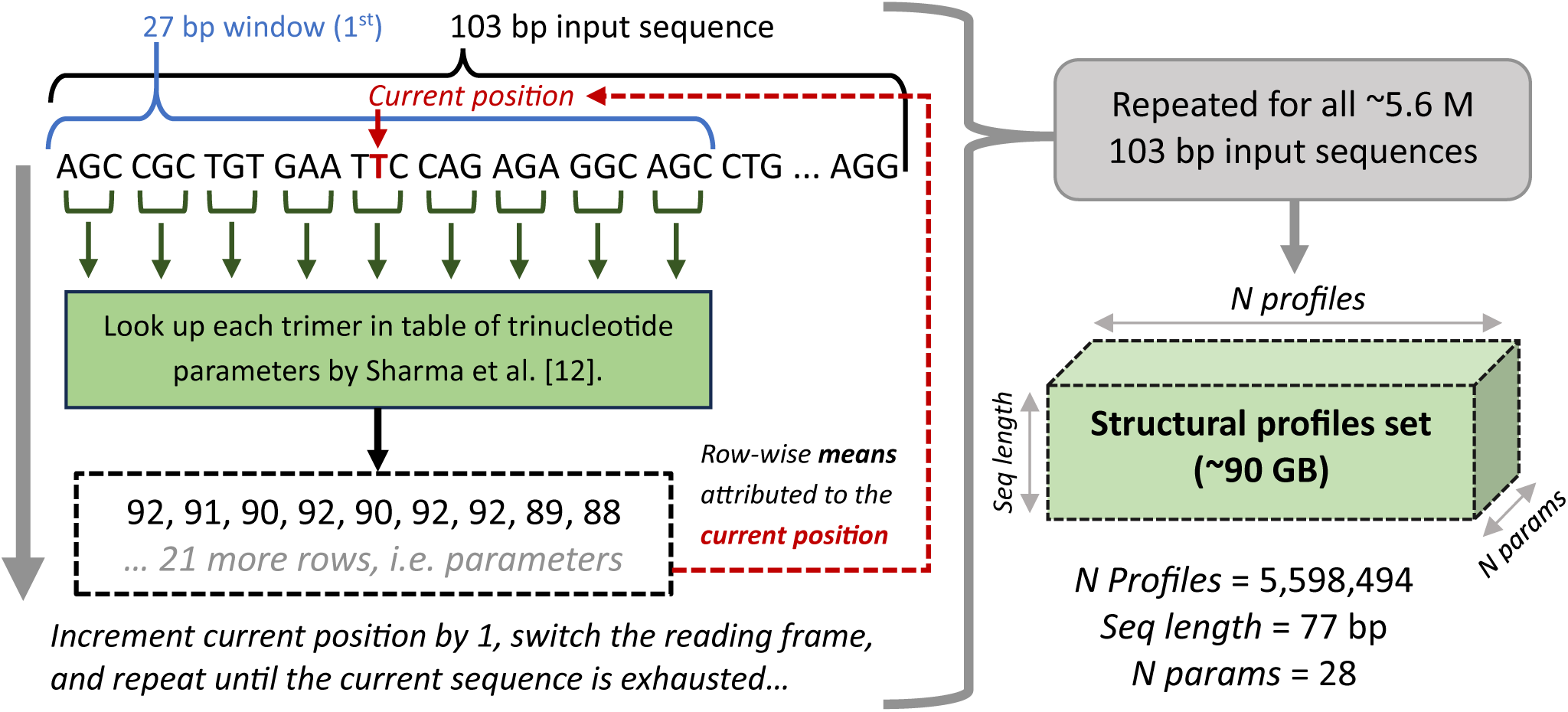
Visual summary of the structural profile generation algorithm. The steps on the left are simplified and omit the use of each trimer reading frame in the mean for each position and parameter. The same steps are also conducted for tetramers to calculate moving averages for 6 additional parameters, which is why the number of parameters is 28 in the final data set represented below.

After sequences were sampled and their structural profiles generated (see **Section 2.3**), the source coordinates of all profiles were ordered sequentially by their position within each chromosome. A random fifth of the coordinates in each chromosome’s ordered list was selected as the testing set. The start coordinate of the first test profile and end coordinate of the last test profile from each chromosome were saved. This created a training/testing partition that allowed much longer sequences to be drawn from within the testing region of each chromosome without the risk of accidental evaluation on mixed training and testing profiles. This enabled a fair sliding window evaluation of the trained models’ performance, detailed in **Sections 2.5** and **3.3**. The outcome of this sampling strategy is depicted below in ***Figure 2***, along with the training profile counts per class and source.

The sampling strategy summarized in ***Figure 2B*** was intended to address a shortcoming of the recent work by Sharma et al. To train and test their model, they sampled 328,368 exon junction sequences, but only 30,140 control sequences, taken from the centers of exons over 1000 bp long [12]. This means that for every ∼10 positive training cases, ChemEXIN only saw ∼1 negative case, but it is intended to be used on sequences where negative cases will occur much more frequently. In the application of machine learning for classification, class imbalance can pose problems. For example, underrepresentation of minority classes in the training set can lead to a model that does not perform well on them in new input data. This is important to consider; however, for generalizability, it can also be critical that the training data is as representative as possible of the true population, imbalanced or not, that the model will see during inference/prediction. Representationally sampling each class relative to others is not always feasible or practical. For example, to sample intergenic sequences at a comparable ratio to others as they occur within the human genome without under-sampling valuable boundary/minority cases, the size of the data set used in this work would have needed to increase by over 7-fold. The full data set represented in ***Figure 3*** already took up ∼90 GB, so this would have made the training times and storage requirements impractical on the available hardware. One workaround for this type of challenge is using sample weights. Sample weights are used in the loss function and penalize the model more or less intensely for classifying members of each class incorrectly. This approach is not without its caveats and limitations, but it is common practice and, in this case, helped to avoid an unacceptably high false positive rate. The sample weights in ***Figure 2B*** above were estimated based on current high-level knowledge about the human genome, e.g. its size and approximate composition. Together, these sample weights should push each model toward optima that are better aligned with the distribution of classes encountered when moving through the human genome, as well as other genomes, than with the training set distribution.

### 2.3) Structural profile generation

While the primary data source for this work was GRCh38 and its RefSeq MANE Select annotation, the models discussed next were not trained directly on sequence data. Instead, they were trained on the moving average of the per-position values of each of the 28 structural parameters. Nucleotide tri- and tetramers were mapped to their corresponding parameter values using the table from Sharma et al., discussed in **Section 2.1** [12]. While describing the full algorithm is impractical, its general premise is depicted in ***Figure 3***. For this step to be possible in Python on a consumer-grade computer in a reasonable amount of time, the profile generation algorithm had to be parallelized. This was done using the multiprocessing library, namely the *imap()* function, as well as efficient algorithm implementation to reap impactful levels of improvement from multi-threading. The profile generation algorithm skipped processing any sequence containing N’s, in order to avoid trimer/tetramer ambiguity and inaccuracy. The numbers reported in ***Figure 2B*** and ***Figure 3*** reflect the sequences that passed this filter.

After the structural profile set was generated, the profiles reserved for training were separated by source (see ***Figure 2B***) and then averaged at the nucleotide level, essentially corresponding to a flattening of each subset along the “N profiles” axis depicted in ***Figure 3***. Introns and intergenic controls were lumped together in this operation. The average profile for each source/subset was z-normalized using the mean and standard deviation of the training set, which can be seen in ***Figure 6***.

### 2.4) Model selection and implementation

As discussed in the introduction, part of the aim of this work was to use deep learning models for the classification, because they have recently shown promise compared to previous state-of-the art *ab initio* gene detection methods. Keras 3 is a deep-learning application programming interface (API) for Python. It was chosen for model implementation in this work primarily because of its low learning curve. TensorFlow was used as the backend, which facilitated the use of GPUs for model training and validation. This was critical for the viability of the training, as doing these steps on a CPU can take orders of magnitude more time.

Three model architectures were selected and evaluated here. The first of these had a single bidirectional LSTM layer (BiLSTM), followed by a hidden dense layer with rectified linear unit (ReLU) activation, and then a final dense layer with softmax activation. This model will be referred to as “BiLSTM” (see ***Figure 4A***). It was chosen because recurrent neural networks (RNNs), such as LSTMs, are considered well-suited for sequence and time series classification problems. “Bidirectional” means the LSTM layer reads the input sequence in both directions, which can be beneficial and is acceptable for this application. The next model involved a temporal convolutional network (TCN), followed by a hidden dense layer with ReLU activation, and then a final dense layer with softmax activation. This model will be referred to as “TCN” (see ***Figure 4B***). An implementation of the TCN architecture, available as a package on GitHub called “keras-tcn”, was used in this model [20]. The authors of that package also authored an arXiv preprint, which showed that this architecture can outperform recurrent networks like LSTMs on standard benchmarking data sets for sequential problems [21]. Perhaps just as importantly, the TCN is more memory efficient and notably faster to train than RNNs [21]. That claim was strongly supported by the training times observed for all 3 models (see supplemental data in **Section 5.1**). These types of performance benefits can have a notable impact when using consumer-grade computers with limited random-access memory (RAM). The third model uses a combination of convolutional layers and BiLSTM layers to extract features from the input across different scales. This is a form of multi-scale dense attention architecture that the authors named MBDA-Net. This architecture was implemented using Keras and adapted for sequence classification, although the authors designed it for hyperspectral image classification [22]. The original architecture of MBDA-Net can be seen in ***Figure 1*** of that paper, and modifications were mainly made to the input and output layers. As such, this model will be referred to as “MBDA-Net” (see ***Figure 4C***). Like Helixer and Tiberius, MBDA-Net utilizes both CNN and BiLSTM layers [4,6,22]. All three models had an input shape of 77 by 28, i.e., profile length by number of parameters. All models had an output shape of 3, i.e., a probability of the input profile belonging to each of the classes: intron-exon, exon-intron, or control.

**Figure 4:**
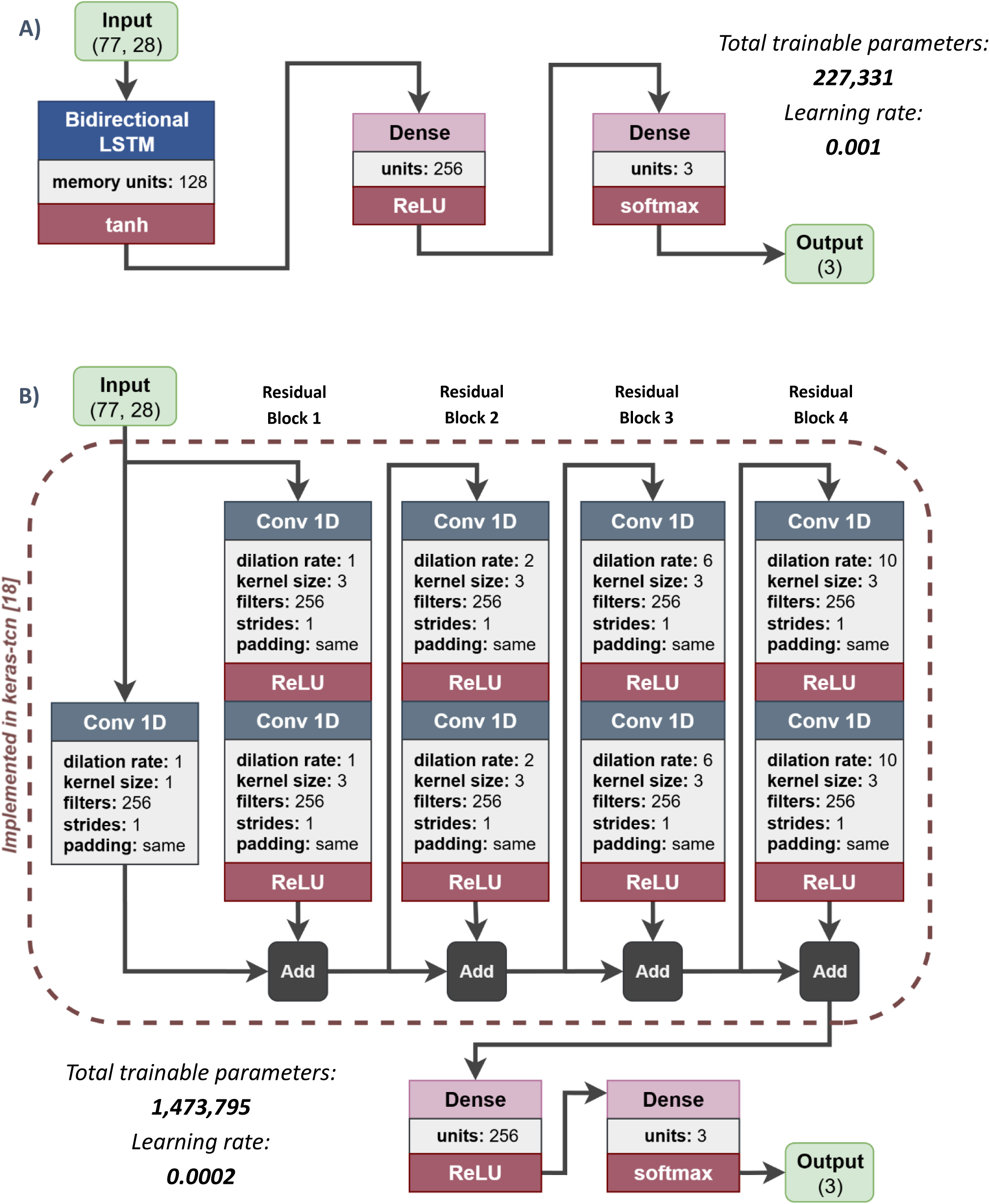
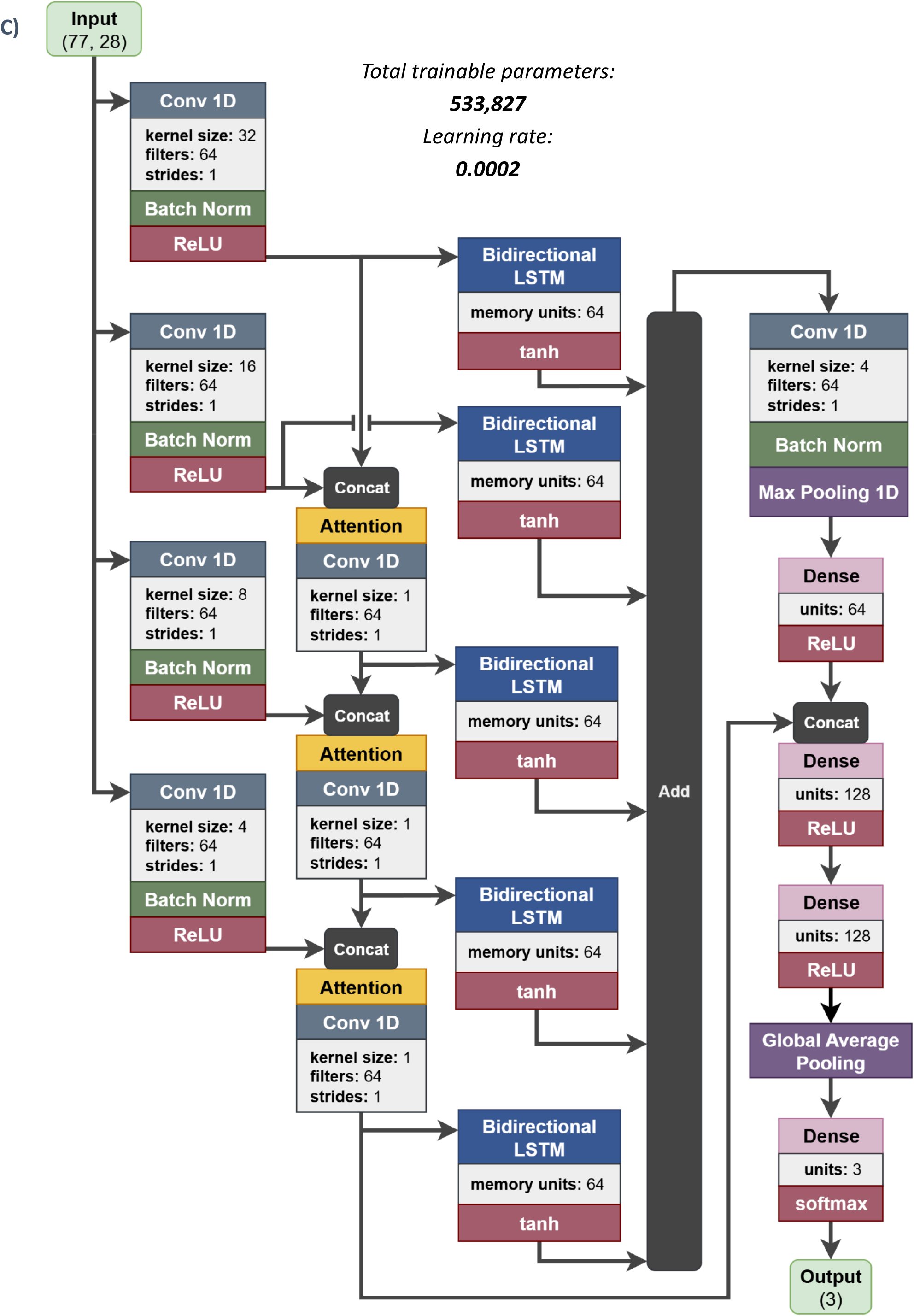
Final model architectures that were trained and evaluated, with key hyperparameters noted for each layer. The total number of trainable hyperparameters and the learning rate used during training are also noted near each model in italicized font. Keras defaults can be assumed for all un-defined hyperparameters in each layer. **A)** BiLSTM-based model. **B)** TCN-based model with a receptive field size of 77. The parts of this model that were implemented in the keras-tcn package are within the dashed outline [20, 21]. A padding type of “same” means the convolutional layer is non-causal. The convolutional layer with no activation function is used to convert the input data into a form that is suitable for addition with the output of the first residual block [21]. **C)** MBDA-Net-based model (see next page). This is an adaptation of a model by Gao et al. [22].

Because of time constraints and compatibility issues with Keras Tuner, a rigorous hyperparameter search was not conducted for these models. Instead, model hyperparameters were chosen based on practical considerations like model size, training time, and epochs until convergence. Initial hyperparameters were chosen for each model via educated guesses based on the complexity of the problem and information from the authors of keras-tcn and MBDA-Net [20–22]. A few critical hyperparameters were then improved upon for each model by limited-epoch trial-and-error runs in a small subset of the training data. This is far from an ideal optimization, but as will be seen, the resulting training and testing metrics were satisfactory. The TCN hyperparameters were largely determined by matching the receptive field of the model to the largest input sequence length (here 77 bp), as per the author’s suggestion [20]. ***Figure 4*** provides an overview of the 3 model architectures and final hyperparameters used during training to generate the results discussed later. As seen in that ***Figure 4***, BiLSTM was a smaller and less complex model than either of the other two.

### 2.5) Model training and evaluation

All 3 models were trained using batches of 64 structural profiles per iteration, with a maximum of 10 epochs allowed. Training was stopped when the validation loss (categorical crossentropy) failed to improve by over 0.001. During the hyperparameter selection process, these were observed to be reasonable choices, and no model ever failed to converge before 10 epochs at any point. Sample weights were used in the loss function that were based on the source of each profile, rather than the class. These were shown in ***Figure 2B***, and the rationale behind them was discussed at the end of **Section 2.2**. As mentioned earlier, the models were trained on 80% of all the profiles generated, and tested on 20% of them. This train/test partition was done in a way that approximately maintained the source and class proportions shown in ***Figure 2B*** in both the training and testing sets.

Normalization was conducted on each profile before any model received it for training or testing. The normalization process involved two steps. The first was z-normalization, in order to center each parameter profile around the global training mean, shift it to 0, and put its deviations from the mean in either direction in terms of the global training standard deviation. After that, min-max normalization was conducted on the z-normalized profile to put all values between 0 and 1. The mean and standard deviation for each parameter across all 4,478,806 training structural profiles were pre-computed prior to training. The post-z-normalization minimum and maximum values for each parameter across all training profiles were then also pre-computed. This step required batch-wise loading of profiles and parallelization to be possible with less than 16 GB of available RAM and feasible in a timely manner. The pre-computed normalization parameters extracted from the training data were also used to normalize the testing profiles, as well as any other profiles used for inference later on.

Because the training and testing data sets did not fit into available RAM, a data generator class was required to prepare the training and testing data batch-wise [23]. During structural profile generation, individual profiles were appended to a single NPY file using NpyAppendArray [24]. During training and validation, data was accessed and prepared batch-wise from the data file using NumPy’s memory map functionality. This also alleviated file read time as a training bottleneck.

All training and testing metrics shown and discussed in the next few sections were computed using the Keras 3 metric class system. Built-in metric classes were used wherever possible, e.g. for calculating precision and recall for each class across a range of thresholds. That said, custom metric classes were needed for calculating the per-class false positive rates (FPR), as well as the per-class areas under the Precision versus Recall (PR) and Receiver Operating Characteristic (ROC) curves. All metrics from each epoch’s training and validation phases were recorded in a CSV file using a Keras callback. Early stopping when the loss failed to improve significantly was also managed with a callback, as was the saving of model weights from the epoch with the lowest validation loss.

Given the crude hyperparameter tuning described in the previous section, 5-fold cross-validation was conducted in the training set to get a better sense of how the selected models might generalize before proceeding with the final training and test set evaluation. Please see the supplemental results in **Section 5** for the training times and cross-validation results.

After training on the full training set and subsequent testing in the full held-out set, a third form of evaluation was conducted that is more representative of these models’ intended use case, i.e., on sequences/profiles longer than the 77 bp input window. Three sets of sequences of sequences were sampled from the regions of the chromosomes reserved for testing (see ***Figure 2A***). The sample sets consisted of 10000 sequences of length 601 bp, 1000 sequences of length 6001 bp, and 100 sequences of length 60001. Each set included only unique, non-overlapping sequences of the chosen size. These were sampled by randomly selecting the source coordinates of 77 bp profiles from the testing set, expanding the coordinates in either direction to the desired length, checking for overlap with previously sampled sequences on the same strand, and retrieving the new sequence if it was unique. These steps were repeated until the desired number of sequences was retrieved. Using each of these sequence sets to evaluate a model corresponded to the model making inferences for a total of ∼6 Mb. As such, these sets were chosen to test the effect of increasing sequence length on model performance. It would have been informative to test a few much longer sequences, but this required notable adaptation of the model post-processing pipeline due to RAM limitations, and there was not enough time. For each sample set, each of the models was fed the normalized structural profile of each sequence in sliding 77 bp chunks (see next section). The model’s per-nucleotide probabilities for each of the 3 classes were then fed into the post-processing pipeline, which translated boundary/nucleotide-level predictions into exon-level predictions. Please see the next section for details on post-processing. Based on these final exon predictions for all the sequences in a set, the performance metrics shown in ***Figures 8 - 10*** were calculated by counting true positives and false positives at the boundary and exon levels, as well as counting the total number of model predictions, and total number of true exons (mapped from the RefSeq MANE Select annotation). The boundary and exon-level metrics shown in ***Figures 8 - 10*** were based on strict true and false negative counts. For example, at the boundary level, a predicted exon start had to match the reference annotation precisely in both position and type of boundary to be counted as true positive. At the exon level, an exon had to match exactly in start *and* end position with the truth/annotation. This is not the only acceptable way to evaluate performance at the exon-level, and more forgiving methods exist [1]. The boundary and exon-level performance for each model in these sequence sets were tallied and evaluated for a range of exon-level acceptance thresholds from 0.05 to 0.95 in increments of 0.05. Please see the next section for an explanation of that threshold.

### 2.6) Model post-processing

As covered in the introduction, *ab initio* gene prediction is a challenging goal. The raw nucleotide-level predictions of even a well-formulated and well-trained model would not be very useful as the final output. Exon-level predictions are the bare minimum, and gene-level predictions are preferable. To accurately achieve that level of output, state-of-the-art tools and models have robust and complex post-processing capabilities, which may or may not be embedded in the classifier’s trained architecture [2–6]. While *ab initio* feature detection with gene-level prediction can take years to develop, some level of basic post-processing was needed to somewhat reliably convert the trained models’ predictions into useful output and simulate a more realistic use case.

The aim and approximate steps of the post-processing pipeline are represented in the panels of ***Figure 5***. First, nucleotide-level predictions (shown in ***Figure 5A***) are obtained by taking the input sequence, generating its structural profile (see **Section 2.3**), normalizing it using the population statistics from the training set (see **Section 2.5**), splitting it into as many “sliding” 77 bp windows as possible (each offset by 1 bp), performing model inference for each window, and aggregating the class probabilities back into their respective positions in the overall sequence. The 77 bp input window of the model was centered on features during training, so predictions correspond to the central nucleotide of the window. Next, the nucleotide-level probabilities are converted into predictions by an algorithm that checks each position represented in ***Figure 5A*** and calls an exon start or end if p > 0.2 for either class. It prioritizes starts if the probabilities of both classes exceed this threshold at a position. Otherwise, all positions are assumed to be controls. Calling the class with the maximum probability at a nucleotide (i.e. argmax) was not chosen for this step because slightly better performance was observed in the test set during training and cross-validation using per-class thresholds. The specific threshold of 0.2 was chosen based on the cross-validation results in the training set, as it seemed to strike a decent balance between precision and recall for the boundary classes without allowing the false positive rate to get too high (see **Section 5.2**). Also, this pipeline’s aim was to avoid being too restrictive on boundary-level predictions at the beginning, so that there was a good supply of exons to filter out via exon-level criteria. In theory, exon-level filtering should have the potential for higher precision and accuracy because boundary-level predictions can be considered within their genomic context.

**Figure 5:**
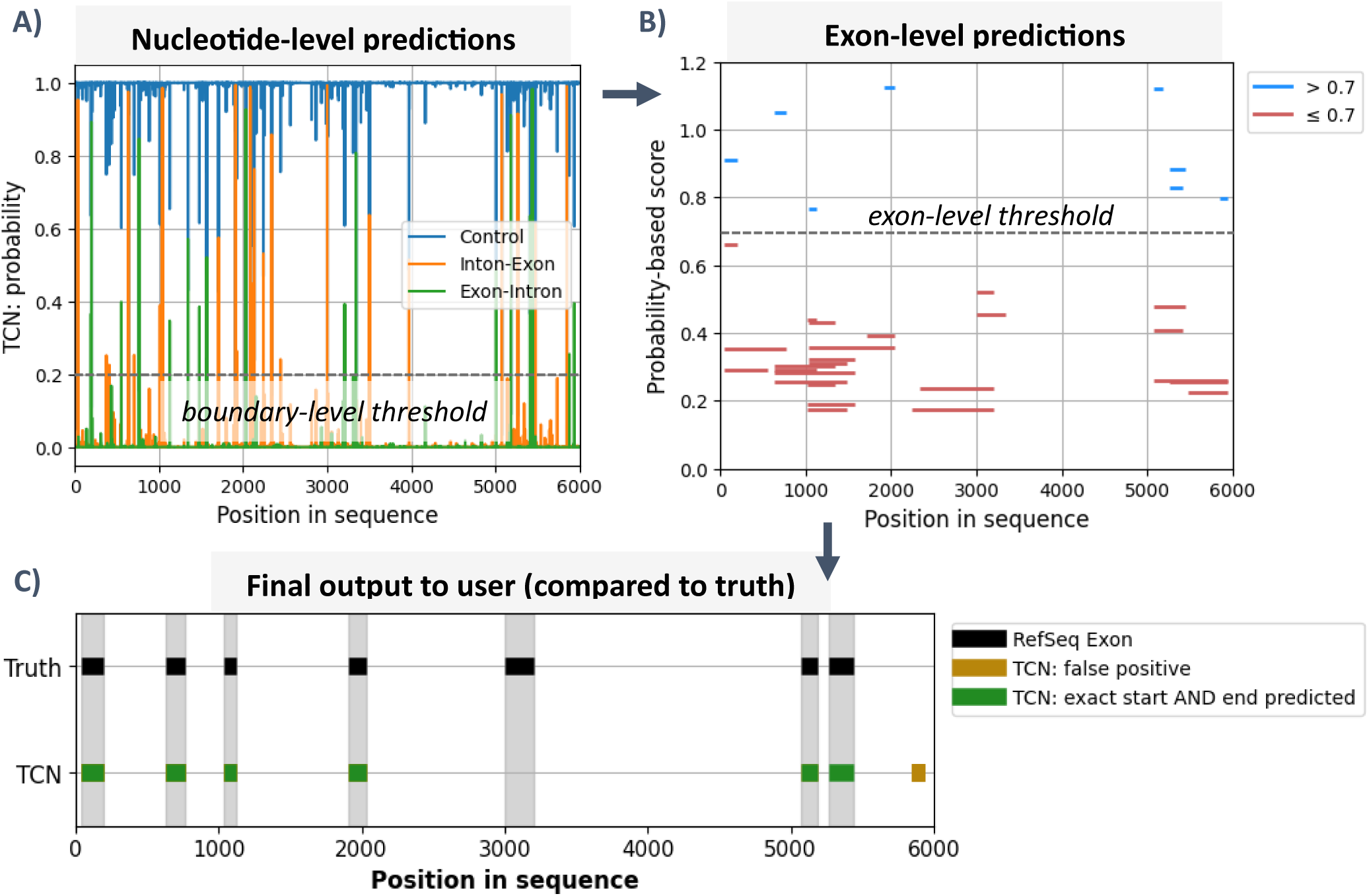
An illustration of the aim and purpose of the post-processing pipeline, which is to take the **A)** raw nucleotide-level predictions from the trained model and translate them into **B)** the possible exon-level predictions, which are ranked and filtered to create a final list of predictions for the user that are (hopefully) likely to be true on average. All plots below correspond to real steps in the post-processing pipeline for the same example sequence: NC_000016.10:15714348-15720348(-). The use of thresholds is a bit oversimplified below, but this figure still generally reflects where and how they come into play (see main text).

The threshold-based boundary predictions outlined above lead to a list of many possible exons, depicted in ***Figure 5B***. These need to be filtered to produce the final exon-level predictions shown in ***Figure 5C***. This step is crucial and can make or break the model’s utility. More advanced approaches would consider the likelihood of *combinations* of exons, given information about typical gene structure, feature length distributions, and biological rules in that type of organism, e.g. via a trained HMM [2–6]. Making a gene-level prediction pipeline was beyond the scope and allotted time of this work, although it is discussed in **Section 4**. A crude but analogous stand-in for more advanced approaches is to rank the exons predictions by their approximate probability within the genome. This was done by first taking the mean probability of each predicted exon’s start and end probabilities, in order to factor-in the extent to which the start and end were high-confidence predictions from the model. The probability of each exon-level prediction was then divided by the absolute value of the log_10_ of the probability of that prediction’s length. Probability as a function of length was estimated via interpolation from a data set on exon frequency versus length in the human genome, discussed in **Section 2.1** [18]. Predicted exons greater than 1000 bp were filtered out because the data set did not include lengths past that point. This re-ranking step is already done for the exon predictions seen in ***Figure 5B***, which is why near-median-length exons are enriched at the higher scores. To go from ***Figure 5B*** to ***5C***, a greedy clustering algorithm, which takes the adjusted exon-level probability score into account, is first is used to enforce the rule that a maximum of 1 exon prediction can exist for any stretch of sequence covered by multiple predictions (not shown). After the predictions are cleaned up in this way, an exon-level score threshold is used to accept/reject exon predictions, which is depicted via color-coding in ***Figure 5B***. The greedy clustering algorithm ensures that there are far fewer predictions going into the threshold/cutoff step by removing conflicting/redundant exons, which matters for much messier intermediate prediction sets than the one seen in ***Figure 5B***. Exon-level filtering then produces the results seen in ***Figure 5C***. The specific threshold chosen influences the precision-recall tradeoff, which is why a range of values were tested in the sliding window evaluations (see ***Figures 8 - 10***).

The post-processing pipeline outlined above was developed by tinkering with a few hundred 4001 bp sequences. These were obtained via random sampling from the training set coordinates and then expanding the coordinate windows to 4001 bp. This post-processing pipeline is far from optimal and relies on approximate high-level information about the human genome. It sacrifices the potential of reaching a high recall for the potential of accurately predicting a specific subset of all exons, i.e., those near the median length. The strengths and limitations of this approach, as well as its validity within the context of *ab initio* gene prediction, are discussed in **Sections 3** and **4**.

### 2.7) Use of ChemEXIN

The performance of BiLSTM, TCN, or MBDA-Net, coupled with the post-processing pipeline, was compared to that of ChemEXIN in sets of sequences sampled to simulate more realistic use cases. Please see the end of **Section 2.5** for information on the sequences themselves and **Section 3.4** for the comparison results. ChemEXIN was released on GitHub as an open-source command-line tool [12,25]. As is, it appears to process only one sequence at time, and user input is required to select which model to use (one trained on human, rat, or worm data) and which seeding threshold to impose for predictions (0.70, 0.80, or 0.85). This is not conducive to generating automated predictions for large sets of sequences. As such, to facilitate comparison with the other models, the source code for ChemEXIN had to be modified. Care was taken to make modifications only directly related to the tool’s ability to process numerous sequences in an automated fashion, and not to change the behavior intended by the authors in any way that could affect model performance. The changes mainly involved alterations to “main.py” so that it could loop through all sequences in a FASTA file and make predictions for them, each time using the model trained on human data and a seeding threshold of 0.85 without asking the user. Most print statements in all modules/scripts were also commented-out for speed, because calls to *print()* add notable time over thousands of iterations. Also, the *prediction()* function definition was moved from “run_model.py” into “main.py” and altered so that the model was only loaded once at the beginning, rather than for every iteration/sequence. This was done for optimal TensorFlow speed during inference, but it did not change the predictions in any way compared to those from the original source code.

## 3) Results

### 3.1) The average profiles

The observation that motivated this work can be seen in ***Figure 6***. Mishra et al. and Sharma et al. included this type of plot in their papers, and it provides an impression of what happens around the intron-exon and exon-intron boundaries on average compared to profiles from other locations [7,12]. ***Figure 6*** illustrates that something conformationally distinctive is predicted to occur around the boundary sites. The variation in the profiles sampled from within exons, introns, and intergenic regions is random such that the mean profile of all profiles from a given source appears relatively flat once scaled by the standard deviation of the training set. Notable variation does occur in these sets of profiles; however, the random variation of all the individual profiles effectively cancels out when they are averaged and normalized. Also, the lines for exons and introns in ***Figure 6*** are noisy, but appear perfectly flat because of a combination of the z-normalization and the axis scale. Meanwhile, the profiles for sequences around the junctions between introns and exons tend to co-vary in certain directions and at certain relative positions on average. This covariation is still visible in ***Figure 6***, even after scaling by the standard deviation of all profiles in the training set, although it is not very extreme.

**Figure 6:**
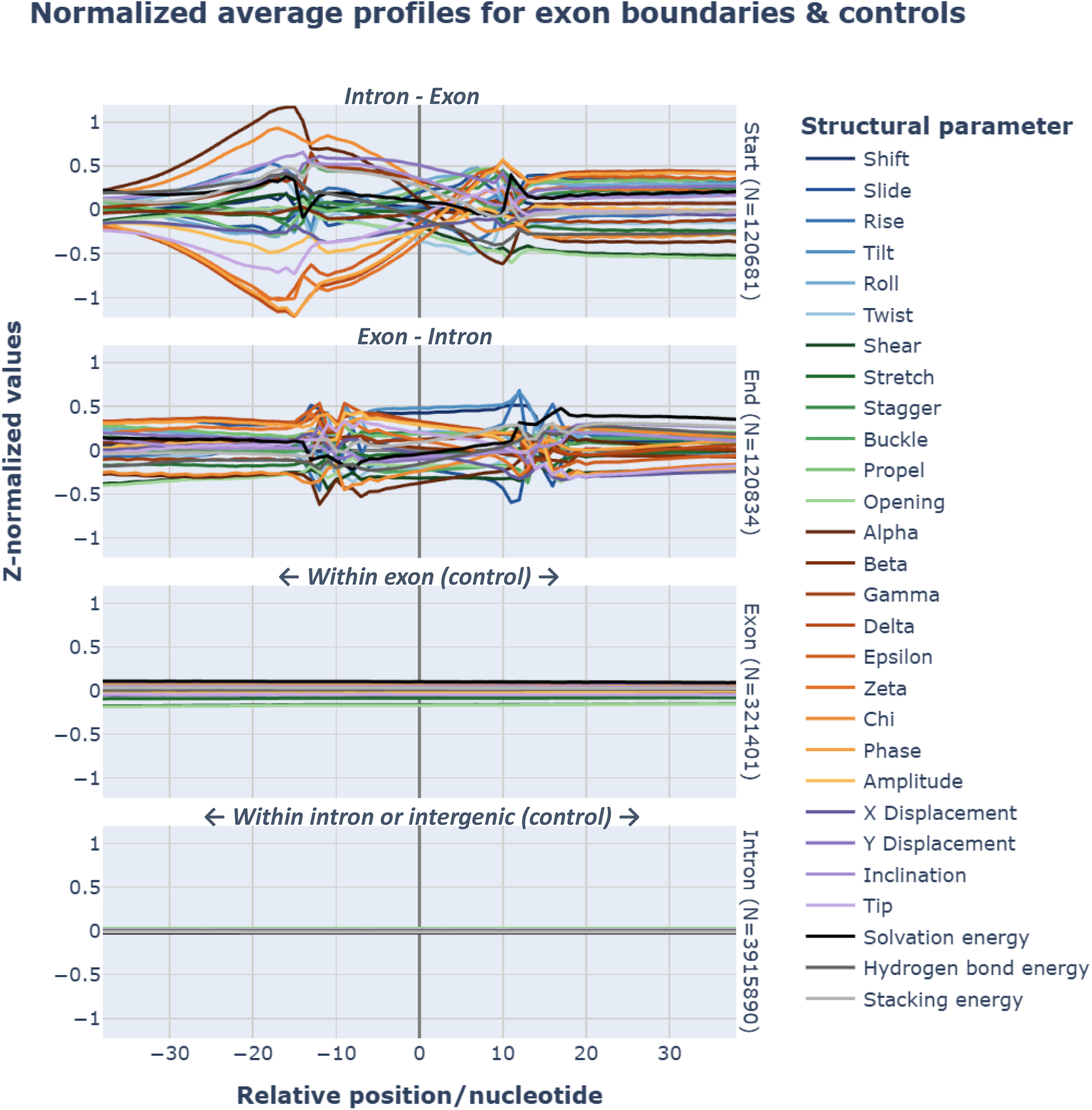
All 77 bp training profiles of a given sequence type were z-normalized using the respective training means and standard deviations (for each parameter) and averaged position-wise. On the x-axis, 0 represents the center position of the source sequences. “Start” refers to the intron-exon junction at the beginning of an exon, while “end” refers to the exon-intron junction at the end of an exon. The “intron” category includes both introns and intergenic profiles, which were lumped together for visual compactness. These parameters have been grouped into five families below, indicated by hue: inter-base pair step (blues), intra-base pair (greens), backbone torsions and sugar pucker (browns/oranges), helical axis (violets), and thermodynamic stability (blacks/grays).

A useful analogy for interpreting ***Figure 6*** is the time-step wise averaging of many sine waves over a specific time interval. If the amplitude, frequency, period, and phase were randomly chosen for each of the wave functions, then the line created by averaging them time-step wise would be noisy, but relatively flat compared to the original sine waves, especially after z-normalization. This is because of destructive interference, coupled with the law of large numbers and the central limit theorem. Alternatively, if only the amplitude of the sine waves was random, and the other parameters were constant across all waves, the time-step wise average would resemble a sine wave. The former case is directly analogous to what occurs when averaging the profiles from exons and introns, while the latter is comparable to what occurs when averaging the boundary profiles.

***Figure 6*** was originally made for a much wider sequence/profile length (401 bp), which is partly what motivated choosing a 77 bp centered training window for the models. As seen in ***Figure 6***, most of the deviation for both boundary types seems to occur within that window, or perhaps an even narrower one. Because these are only average profiles, a wider window was chosen in order to err on the side of not missing any potentially useful signals. Because the average profiles in ***Figure 6*** are normalized, their minimum and maximum values are somewhat informative. They show that, on average, these boundary signals do not deviate much farther than 0.5 to 1 standard deviations from the mean of the profiles of every type of profile sampled. This might imply that the signal-to-noise ratio and class separability pose challenges for the profiles of interest; however, as discussed above, ***Figure 6*** only shows the average behavior of populations of structural profiles. The individual profiles are not homogenous.

### 3.2) Test set performance

***Figure 7*** provides a comprehensive overview of the performance of BiLSTM, TCN, and MBDA-Net on the test set. All models exhibited very similar and satisfactory performance. Minor differences are most apparent when looking at the F1 scores as a function of decision threshold. That TCN and MBDA-Net did not perform significantly differently than the much smaller and simpler BiLSTM model implies that their added size/complexity may not have been necessary to learn the essential features required to distinguish boundary profiles from others, barring a major failure in their hyperparameter tuning. That said, the BiLSTM model had the most variability in its precision-recall performance during cross-validation in the training set (see supplemental Figure 13 in ***Section 5.2***). While not a guarantee of improved generalizability, this may indicate that the other two architectures are more robust.

**Figure 7:**
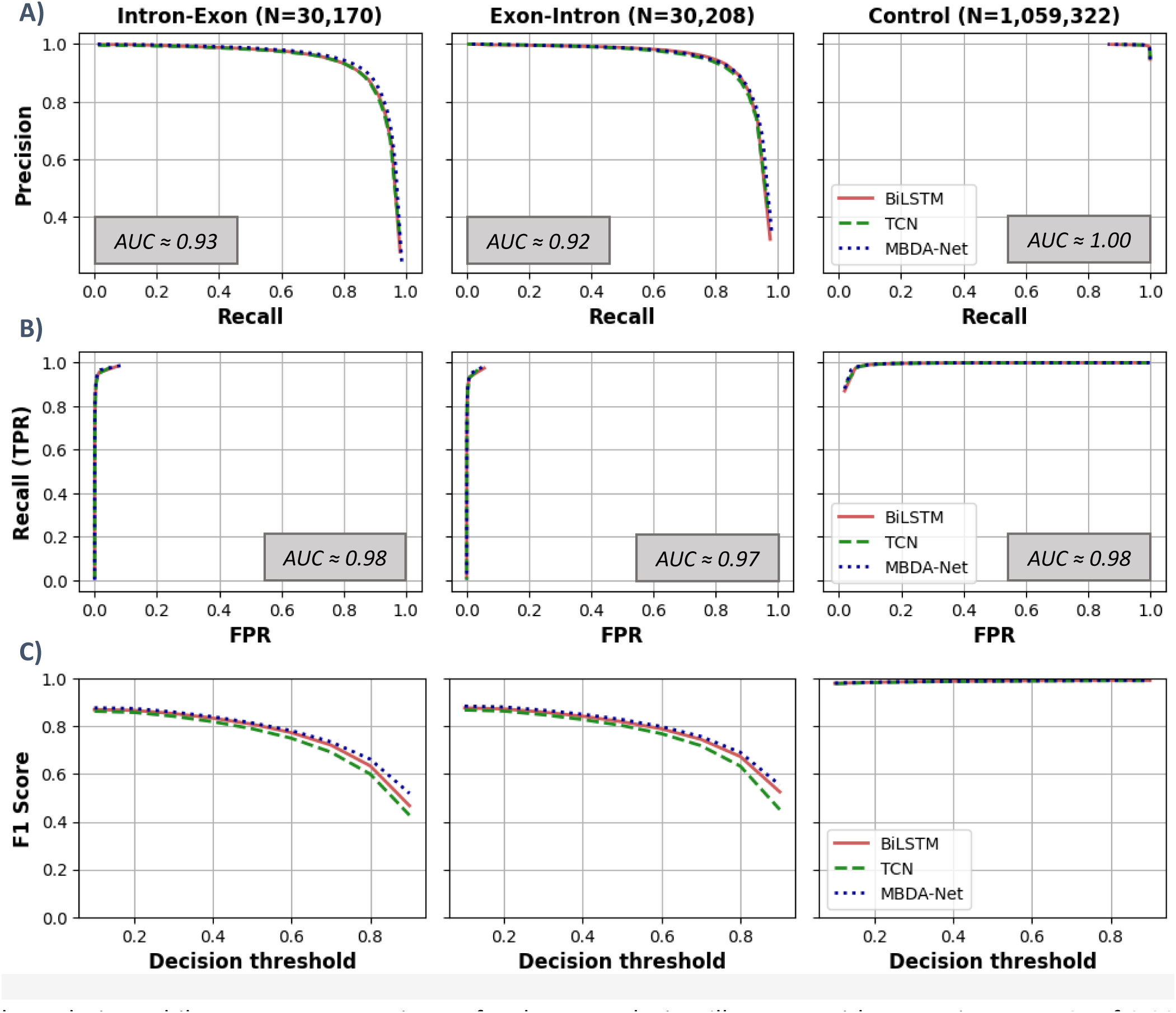
Per-class performance summary in the set of held-out/evaluation profiles. Each column of plots corresponds to the class noted at the top, with the number of that class in the test set in parentheses. Row **A)** contains the precision versus recall (PR) curves. Row **B)** contains the true positive rate (TPR) versus false positive rate (FPR) curves, i.e. the ROC curves. Row **C)** contains curves of the F1 Score, i.e. the harmonic mean of precision and recall, at each of the decision thresholds used in the earlier two rows of plots. Some curves appear incomplete in the first two rows due to a limited number of thresholds being used, but they still capture the performance clearly. The approximate area under the curves (AUC) is noted for the PR curve and ROC surve (mean of the 3 models’ actual AUC values in each case).

Because the 3 models exhibited very similar testing performance, the trends for all of them in ***Figure 7*** will be discussed as one. In terms of precision versus recall, these models performed the best on non-boundary (i.e., control/negative) cases, with an AUC very close to 1. This comes as no surprise, given the much larger quantity of control profiles these models saw during draining. This result shows that the aim of the profile sampling and sample weighting rationale, discussed at the end of **Section 2.2**, was a success. These are models that will not struggle too much with misclassifying negative cases if used correctly, which is a strength in the setting of a genome, where exon boundaries (positives) are a relatively rare occurrence. An *ab initio* gene detection algorithm ought to perform well, even when no genes exist in the sequence. The precision-recall (PR) performance for the exon-intron and intron-exon boundaries, while not as strong as it was for the controls, is still strong, with approximate AUCs of 0.93 and 0.92, respectively. Taken together, the PR curve, ROC curve, and F1 score curve indicate that, with the right choice of decision thresholds, for each class any of these 3 classifiers could simultaneously achieve a decent success rate, capture a decent proportion of all cases, and achieve a reasonably low false positive rate. That said, strictly speaking, choices about optimal cutoffs ought to be made based on results in the training set, which they were (see **Sections 2.6** and **5.2**).

Sharma et al. provided a per-class ROC plot with performance for the convolutional network at the heart of ChemEXIN [12]. The comparable plot for the models trained and evaluated here may imply improved generalizability. That said, a high AUC-ROC is not particularly difficult for a model to achieve, nor a guarantee of its success. Sharma et al. reported test set AUCs of only 0.85 for the boundary classes and 0.78 for controls, whereas the models in ***Figure 7B*** had approximate per-class AUCs of ≥ 0.97 [12].

### 3.3) Sliding window evaluation

While the results in ***Figure 7*** are encouraging, they are only necessary for these models to have the possibility of performing well in their intended use case, but do not guarantee it. Each true or false positive tallied to generate ***Figure 7*** arose from the model being shown a single profile for a 77 bp window. For junctions, that window was conveniently centered on the start or end of an exon. The practical use of this model involves sliding its 77 bp input window 1 bp at a time through the profile generated from the input sequence, which could amount to anywhere from hundreds to millions of predictions (or more). Even if representative, an apparently small false positive rate for any class in the test set could amount to a large absolute number of misclassifications in a real use case. Also, the model could easily struggle with generalization if it is presented with more noise/variation throughout a sequence or genome than it was prepared for during training. Put differently, the training and testing scheme outlined so far, while standard practice in machine learning proof-of-concept papers, was formulated based on what is practical, rather than optimal. As such, a more rigorous and representative test is merited to see how these models might perform during their intended task. For this reason, sequences were sampled from within the regions of each chromosome reserved for the test set so that the models could be evaluated in a more realistic use case. 10000, 1000, and 100 sequences with respective lengths of 601, 6001, and 60001 bp were sampled and converted into structural profiles. Within each sample set, the sequences were unique and not part of the models’ training set. Please see **Section 2.5** for the sampling details and rationale for the sliding window test sequences. Also, depending on the context, trained sequence classifiers can require some level of post-processing to convert class probability predictions for each nucleotide into a more useful level of output. Design choices made in the post-processing can improve or harm a capable model’s final output quality. The details and rationale for the post-processing steps that resulted in the model performance shown below are detailed in **Section 2.6**.

***Figure 8*** summarizes the sliding window performance at the boundary-level, i.e. each model’s ability to differentiate intron-exon and exon-intron sites from the rest. Aside from lumping the two boundary classes together, the metrics in that figure are analogous to those in ***Figure 7*** in their interpretation. The performance of all 3 models is again similar, with some slight differences in terms of the maximum recall that can be reached and the associated false positive rates, especially for the 60k bp sequences (compare TCN and MBDA-Net to BiLSTM). Overall, these models struggle considerably more in this simulation of their intended use than they did during the traditional test set evaluation. At the right thresholds in ***Figure 7***, these models achieved a balance of strong precision and recall, with F1 scores ≥ 0.85 for both boundary classes. Meanwhile, the maximum F1 scores in ***Figure 8*** for each model are only in the ≥ 0.5 range. The best these models can do is achieve decent-to-mediocre precision at the severe expense of recall. The specificity (1 - FPR) of these models is strong overall at the boundary level, even in ***Figure 8***. That said, even a well-managed FPR can lead to a high absolute number of false positives in long sequences, which is one reason why post-processing techniques can be crucial (more on this in **Section 4**). The maximum recall these models could achieve in the longest sequences was mediocre at best (roughly 0.6 - 0.7), which came at the expense of precision. Recall is most tightly limited at the shortest sequence length because of the impact of boundaries lost when the exon’s start or end falls outside of the sequence. As discussed in **Section 2.6**, the post-processing pipeline uses probability at the exon-level to accept or reject boundaries. It does not attempt to predict exons that do not have a start and end within the profile window, which is more likely to happen for shorter sequences and result in missed exons. That these models struggle more with recall than precision is also somewhat by design. As discussed in **Section 2.6**, the post-processing algorithm considers pairs of boundaries, i.e. exons, within the context of the probability distribution of exon length in the human genome. This enhances the maximum precision of these models, but it is a crude algorithm, so exons that fall too far from the median length in either direction have virtually no chance of being predicted. The post-processing algorithm is also part of the reason the performance of all models is fairly homogenous (although not exactly). Better alternatives to the current post-processing solution are discussed in **Section 4**.

**Figure 8:**
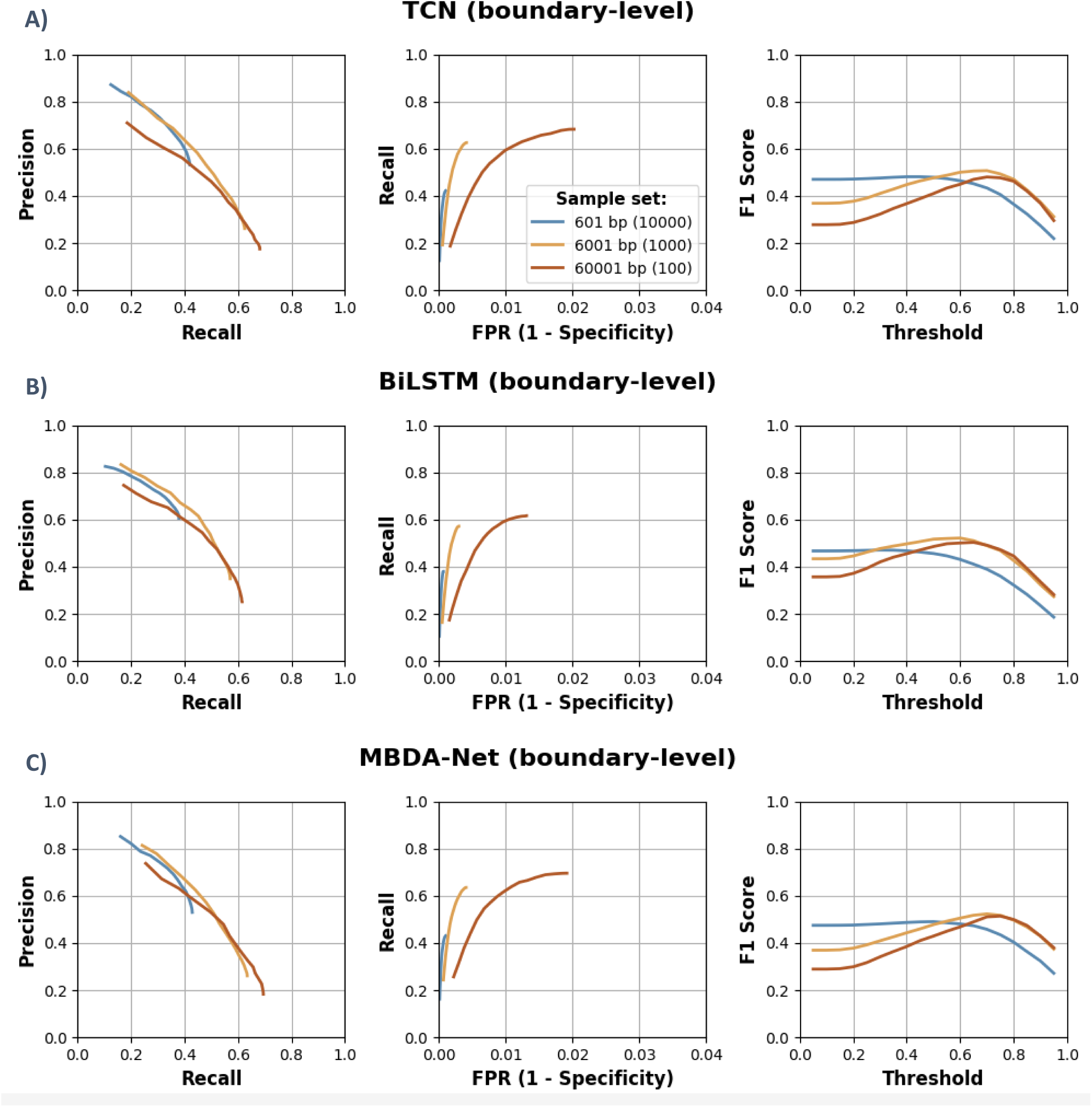
Boundary-level sliding window performance summaries for **A)** the TCN model, **B)** the BiLSTM model, and **C)** the MBDA-Net model in sets of 3 sequence/profile lengths. These results are from models coupled with the post-processing algorithms detailed in Section 2.6. Unique, non-overlapping sequences were randomly drawn from the regions the chromosomes reserved for testing for each sample set/line color (see Section 2.5). The intron-exon and exon-introns performance counts were lumped together to compile these boundary-level results.

***Figure 9*** shows the exon-level performance of these models for the same sliding window sequence sets. Precision and recall were calculated strictly, where only exact matches to the beginning and end of the true exons were counted as true positives. The FPR was calculated more leniently based on the proportion of all non-exon sequence length that was covered by predicted exons. All performance metrics diminish to some extent at the exon-level, which is to be expected because this is a higher bar for each model. Both the precision and recall are even more limited in the PR curves in ***Figure 9*** than they were at the boundary level. This is reflected in the maximum F1 scores, which sit more around 0.45 for exon-level predictions. The false positive rate “explodes” while recall saturates for all models because of the post-processing algorithm described earlier (middle column of ***Figure 9***). There is a narrow subset of all possible model predictions that the algorithm will pass. These are exons with high average probability between start and end sites that fall closer to the median exon length, compared to other overlapping predictions. As such, the correct exons that the post-processing pipeline can predict are enriched at higher exon probability scores. As the final exon acceptance threshold is decreased to allow more exons through, all that happens past a certain point is the addition of low-quality exon predictions that do not overlap well with the true exon positions. The final result is an increasing FPR at a constant recall. This is a tradeoff of the post-processing algorithm that accepts a lower recall overall for a decent precision operating regime (upper left of PR curves in ***Figures 8*** and ***9***).

**Figure 9:**
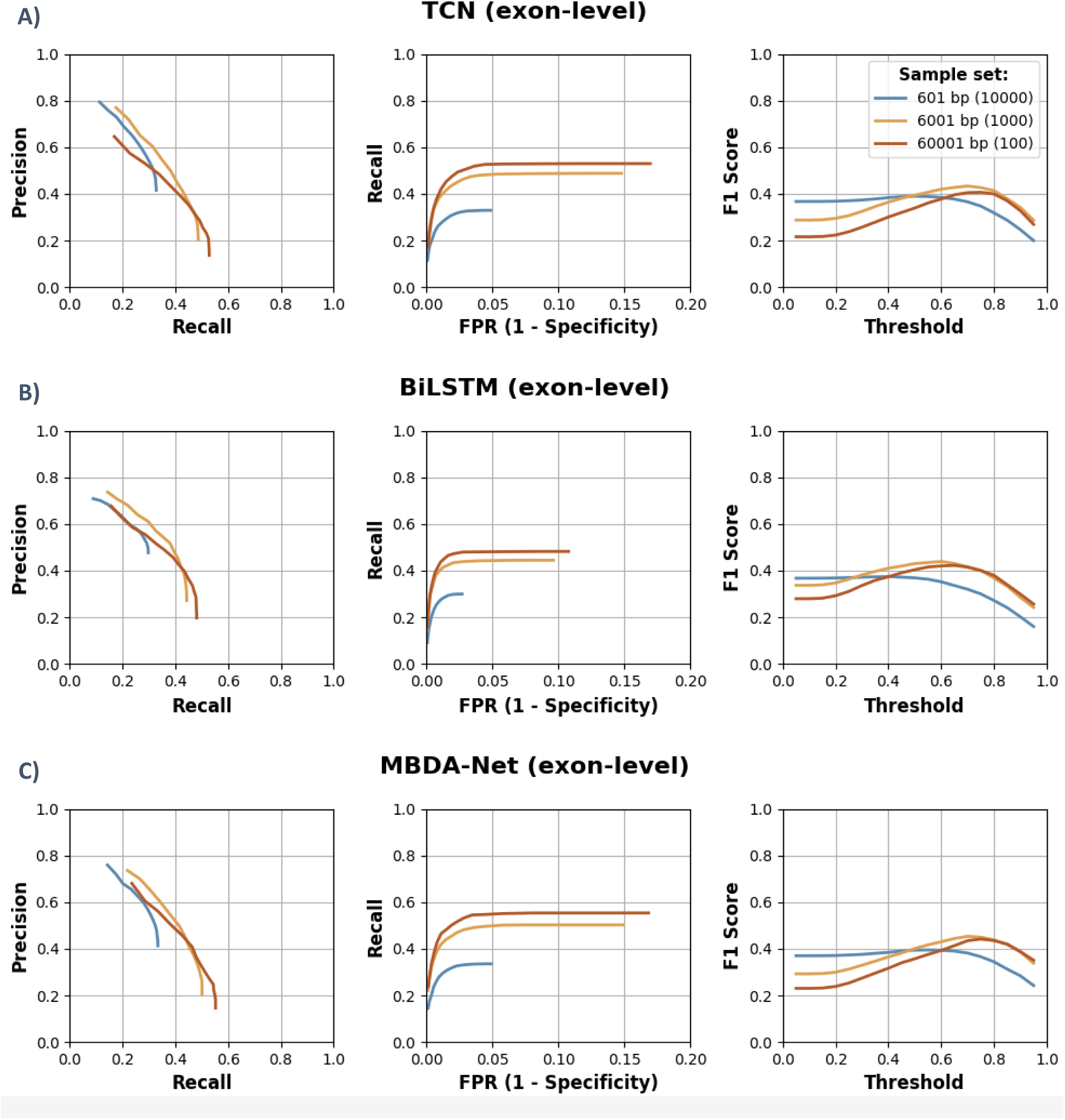
Exon-level sliding window performance summaries for **A)** the TCN model, **B)** the BiLSTM model, and **C)** the MBDA-Net model in sets of 3 sequence/profile lengths. These results are for the same sequences sampled and tested for Figure 8. Precision and recall were calculated based on strict exon successes, i.e. when both the beginning and end exactly matched the truth (RefSeq annotation), rather than predicted exon overlap with the truth. The false positive rate (FPR) metric was calculated more laxly, by considering the proportion of all non-exon length in the sequence/profile that was covered by a model’s exon predictions.

### 3.4) Comparison to ChemEXIN

The plots discussed in the previous sections show models that generalized well in the test set but will probably struggle in their intended use case with the current post-processing strategy. Right now, the best they can do is make predictions with moderate-to-high precision, but only capture a limited proportion of all the exons that exist. That said, how do these models compare to the implementation by the authors of this approach? The title of that paper claims that ChemEXIN has made the detection of exon boundaries “easy”, and includes a table showing impressive performance in 3 species compared to some of the previous state-of-the-art methods, including AUGUSTUS [12].

To explore this question, ChemEXIN was run on the same sets of sequences used to generate ***Figures 8*** and ***9***. While the documentation on GitHub is somewhat unclear [25], the most likely interpretation of ChemEXIN’s output seems to be that it provides windows in which an exon boundary is predicted to exist. Whether an intron-exon or exon-intron junction is predicted within that window appears to be unspecified. As such, the calculation of the performance metrics was adjusted to work with this level of output. Also, ChemEXIN has two types of prediction output, both of which were evaluated and included in ***Figure 10***. For more details on how ChemEXIN was used and evaluated, please see the ***Figure 10*** caption and **Section 2.7**. In ***Figure 10***, the performance was included from ***Figures 8*** and ***9*** for the threshold and model that achieved the most similar recall to ChemEXIN.

**Figure 10:**
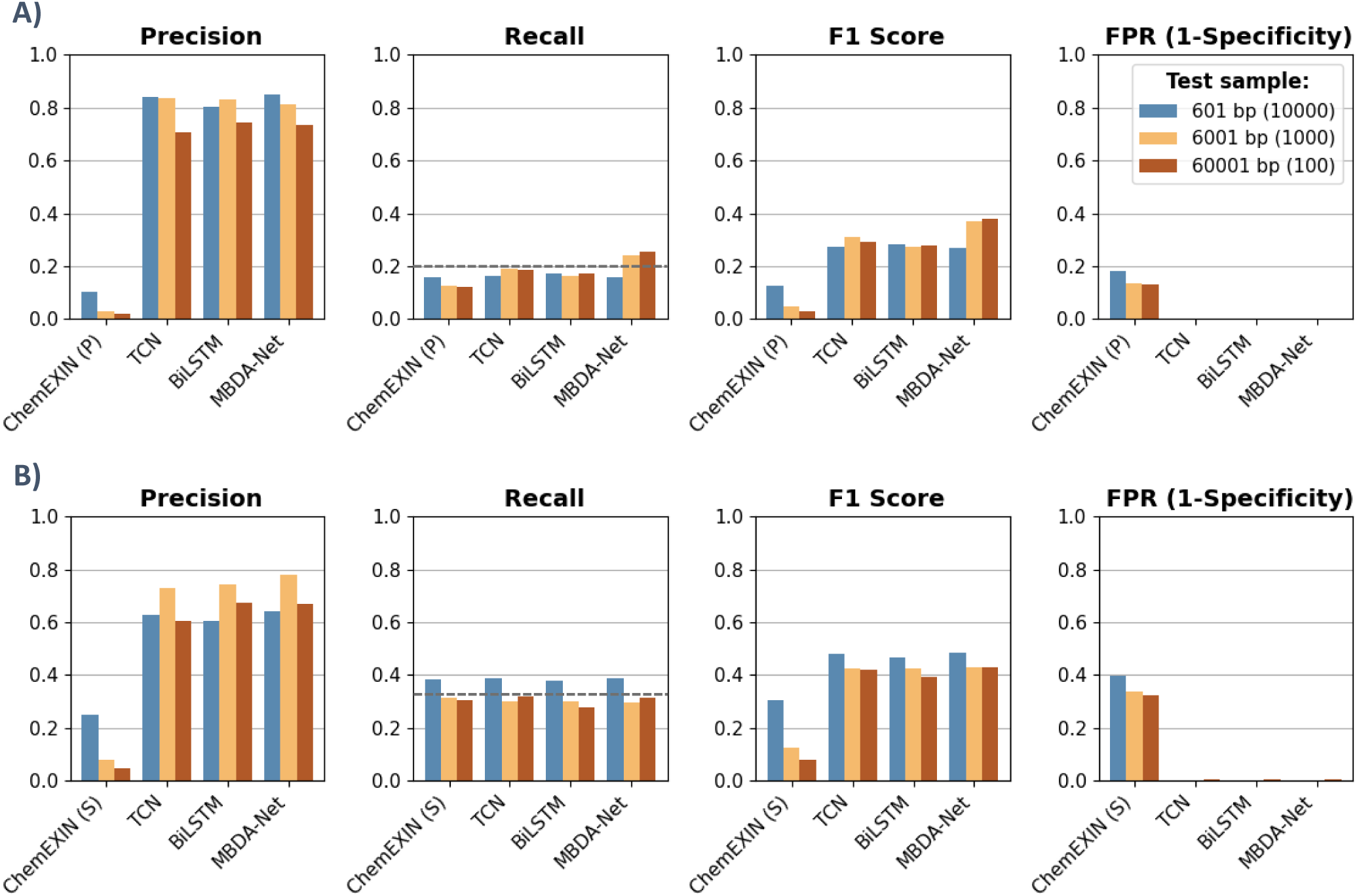
Comparison of the 3 models’ boundary-level sliding window performance to that of ChemEXIN for the same sequence sets at comparable recall (indicated by the dashed lines). **A)** Comparison to ChemEXIN’s primary (P) prediction windows. **B)** Comparison to ChemEXIN’s secondary (S) prediction windows. A seeding threshold of 0.85 was used for ChemEXIN, which was the most precision-leaning option [25]. The threshold varies for the other 3 models to enable comparisons at similar recall (see Figure 8), and it is not directly comparable to ChemEXIN’s threshold, as boundaries are filtered in pairs at the exon-level. ChemEXIN’s precision and recall are based on predicting a window truly containing either type of boundary, while the same metrics for the other models are based on predicting the exact position of an exon start or end correctly. FPR is not visible for BiLSTM, TCN, and MBDA-Net due to scale (see Figure 8). ChemEXIN’s FPR was calculated based on the proportion of non-exon sequence length covered by predictions, just as it was for the other models.

As seen in ***Figure 10***, at comparable recall, either type of prediction from ChemEXIN (primary or secondary) has significantly reduced precision compared to the 3 models discussed earlier. This causes much worse F1 scores compared to those of the other models, which are not great to begin with. The low precision is driven by an unacceptably high false-positive rate. The secondary predictions from ChemEXIN seem to be obtained by expanding the primary predicted window of 50 bp by 30 bp in either direction, based on whether the model predicts other boundaries nearby [25]. That said, the explanation from the authors of why this happens is not completely clear. In either case, the rationale does not change the fact that this expansion of some of the primary predicted windows by a fixed/arbitrary amount, in order to generate a set of secondary windows, increases the recall by non-specifically increasing the odds of hitting any boundary. This is apparent in ***Figure 10***, as the ChemEXIN predictions in ***10B*** increase the recall notably compared to ***10A***, but they also increase the FPR by a comparable amount. Overall, while BiLSTM, TCN, and MBDA-Net do struggle, they are a notable improvement over ChemEXIN’s performance, because they can at least predict the exact location of some subset of the true exons with passable precision. ChemEXIN’s performance is abysmal, which seems discordant with what is reported in the paper [12]. Potential reasons for this apparent discrepancy are discussed at the end.

Finally, while the numbers displayed in ***Figure 10*** are more informative and comprehensive than single cases, looking at how these models are performing within specific sequences adds some context that is hard to communicate otherwise. Visual inspection is important for checking that any algorithm, model, analysis, etc. is behaving consistently with what is thought or assumed. ***Figure 11*** displays the RefSeq MANE Select exons for several sequences compared to post-processed predictions from the MBDA-Net model, as well as ChemEXIN. ChemEXIN’s secondary (S) predictions are shown because they yielded the best performance in ***Figure 10*** (aside from FPR). The seeding threshold used was again 0.85. The predictions of MBDA-Net in ***Figure 11*** were obtained using an exon acceptance threshold of 0.7. Because these are only a few sequences, the results in ***Figure 11*** are not representative of MBDA-Net’s performance seen in ***Figures 8 - 10*** for the corresponding sequence lengths. They were primarily chosen to highlight and discuss specific behaviors of all 3 of the models discussed above, as well as ChemEXIN.

**Figure 11:**
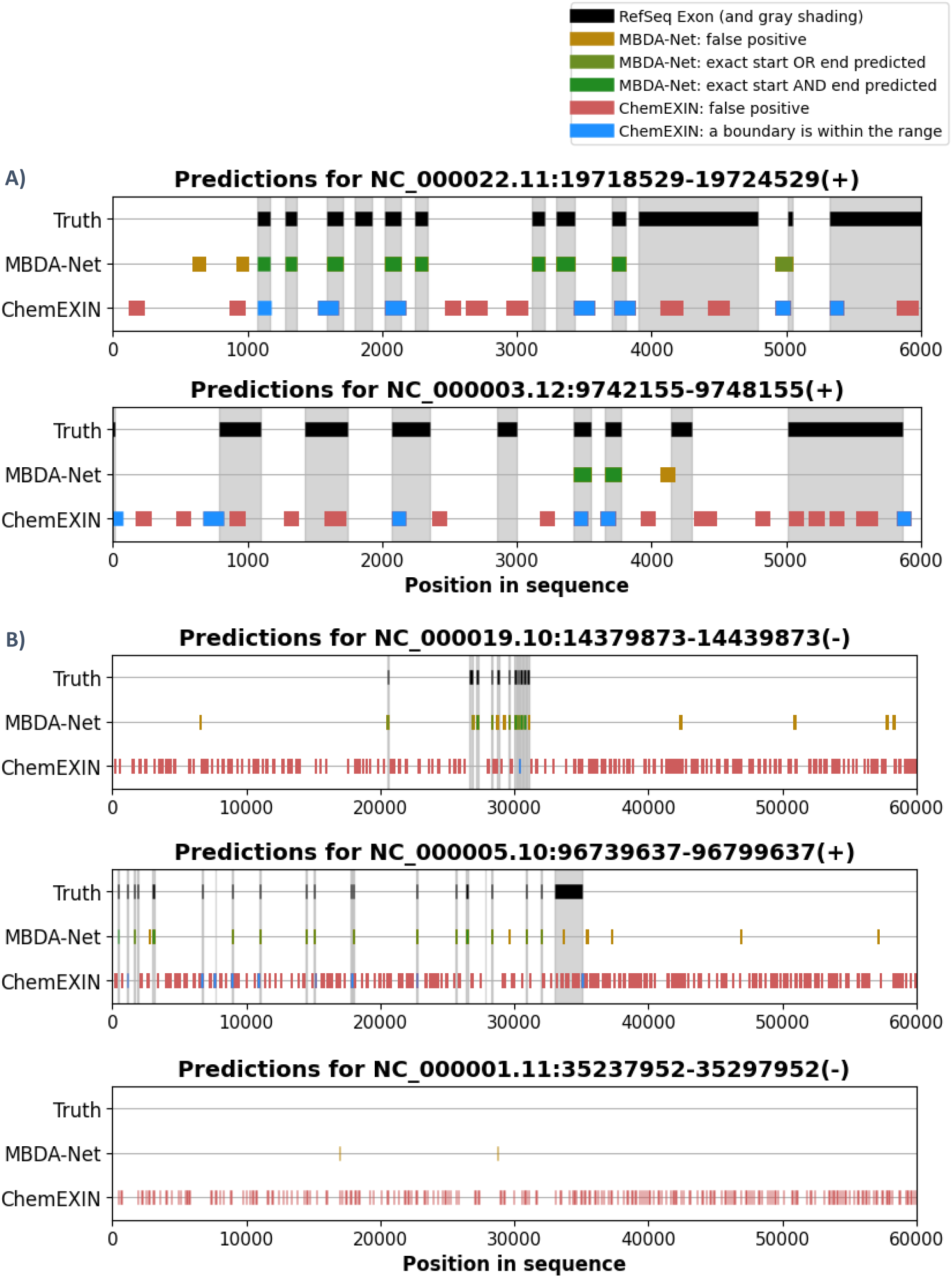
Comparisons of ChemEXIN (S) and MBDA-Net predictions to the true features for **A)** 2 sequences of 6k bp and **B)** 3 sequences of 60k bp. These sequences come from the respective test sets used to evaluate the models at different sequence lengths (see Figures 8 - 10). The genomic coordinates in GRCh38 for each sequence are displayed above each plot. ChemEXIN outputs a window in which it predicts an exon boundary (start or end) to exist, while MBDA-Net aims to predict the exact beginning and end of a full exon.

ChemEXIN’s low precision and high false positive rate are quite apparent in ***Figure 11***. Although MBDA-Net certainly produces false positives where no features exist, this is much more tightly controlled than it is for ChemEXIN. This is largely a result of a more comprehensive training scheme, which exposed the models to many more negative cases drawn from throughout the genome, which factored into the loss function more heavily. The constant stream of ChemEXIN false positives in every sequence, including the one with no features (***Figure 11B***, bottom), shows that it would not make a viable *ab initio* gene finding tool, at least as currently exists on GitHub. Also, the prediction of an approximate location of a boundary, without specification of the type of boundary, is not a very helpful level of output for a user to infer exons from. As noted in the introduction, even the early gene prediction tools often aimed to at least provide exon-level predictions, whether they were reliable. After reviewing roughly a thousand plots like the ones in ***Figure 11***, it was still unclear whether ChemEXIN’s predictions are covering some true boundaries for any reason other than the frequent number of predictions being made, coupled with the (apparently) arbitrary expansion of the prediction window.

While ***Figure 11*** shows that MBDA-Net might have some potential, it also highlights some key weaknesses of the post-processing pipeline that it currently relies on for predictions. It is easy to see in ***Figure 11A*** that MBDA-Net regularly misses the longer exons completely. As discussed earlier and in ***Section 2.6***, this is to be expected. The post-processing pipeline that turns base-wise predictions into exon predictions prioritizes exons based on the likelihood of their length within the human genome, but this was implemented rather crudely because of time constraints. As such, the pipeline will never predict an exon longer than ∼1000 bp, and it is very unlikely to predict exons much longer or shorter than the median exon length, which is roughly 120 bp [18]. That said, the pipeline is the primary reason the multiplicity of combinations of predicted start and end sites from MBDA-Net can be filtered into exon predictions with any accuracy. ***Figure 11*** also shows there are different ways to measure success at the exon level, e.g. by considering the level of overlap/similarity for exon predictions that are not spot-on. This has been common practice when evaluating gene prediction programs in the past [1]. For simplicity, a strict evaluation criterion was chosen to generate ***Figure 9***, in which predictions like the yellow-green one in ***Figure 11A*** counted as a false positive at the exon-level.

It seems fair to wonder whether using the length distribution of exons in the human genome to rank exon predictions from MBDA-Net, or any of the other models, introduces information leakage. If a model is being used to predict the locations of exons or full genes within a sequence, would this information be available for every genome? If not, would such a practice generalize well? Is this not part of the information such a model ought to be helping to uncover, e.g. in a newly sequenced genome? First, the length distributions of exons are somewhat consistent across large swaths of the tree of life. For example, this is fairly consistent across many vertebrate species [26]. Second, as mentioned in the introduction, this type of information has been a crucial part of the success of state-of-the art tools for some time. This includes classics like AGUSTUS, as well as modern ones like Helixer and Tiberius [2,4,6]. The difference is that the length distribution data used in this work was not very detailed, and it was deployed as a rudimentary sorting function after-the-fact (see **Section 2.6**). For AGUSTUS, Helixer, and Tiberius, the multi-faceted and context-dependent nature of the length distributions of exons and other genomic features was modeled/learned via the HMM and/or deep learning parts of those tools as part of their training. These models use and encode representations of numerous types of information beyond just length distributions, all of which relate to the layout of a typical eukaryotic gene. This is how they can predict whole genes, not just individual boundaries or exons, with impressive levels of accuracy.

## 4) Discussion

As they currently exist, the models trained and evaluated for this work are not ready for the challenging task of reliably predicting exons within diverse genomes. This comes as no surprise, as state-of-the-art tools like AUGUSTUS, BRAKER, Helixer, Tiberius, and ANEVO are trained on (or use) data from multiple genomes, evaluated in many others, and distill features from sequence data beyond just those associated with exon junctions. While their performance is highly context-dependent, these tools often achieve exon-level F1 scores of 0.6 - 0.9 in animal species and beyond, with recent deep learning tools often achieving scores toward the upper end of that range [2–6]. While BiLSTM, TCN, and MBDA-Net generalized well to the testing set (max boundary-level F1 ≳ 0.85), these models struggled to maintain the same level of performance when processing full sequences (max boundary-level F1 ≳ 0.5, max exon-level F1 ≈ 0.45). That performance could be further diminished in sequences from organisms only distantly related to humans. That said, these models may generalize better than ones of similar complexity trained on sequence data alone. While the structure of DNA is far less diverse than that of RNA or proteins, the higher number of degrees of freedom in the molecular structure domain allows for distinct sequences to produce similar structures. As such, making predictions based on input from this domain might have the potential for better generalizability across diverse species, assuming the predicted structural signatures are accurate and conserved. Also, despite its other shortcomings, the post-processing algorithm used with the models in this work relied on human exon length distribution data, which may at least generalize fairly well across vertebrate species [26]. An informative next step could be to test the performance of these models, with the current pipeline, on sets of sequences from organisms of increasing genetic distance from humans.

While the main conclusion of this work is that the models trained and evaluated are not competitive alone in their current state, the potential of this approach cannot be definitively discounted. The poorer sliding window performance detailed above may be at least partly attributed to the specifics of the implementation, namely the post-processing methodology. This algorithm aggressively ignores exon-level predictions that do not fall around the median length in order to achieve a more acceptable maximum precision. While near-median-length exons occur most frequently, this still puts a major limit on recall. When developing the post-processing algorithm in sequences taken from the training regions, the intermediate predictions were frequently inspected and compared to the truth. Correct long (or short) exon predictions of only moderate probability were frequently discarded later by the algorithm during the length-based adjustment of their probability score. As such, the pairing of the models evaluated here with more sophisticated post-processing may make them notably more competitive. A reasonable starting point would be an HHM or additional deep learning layer that has been formulated and trained to assign probability scores to sets of predicted exons based on whether their lengths and orientation reflect a configuration that is likely within a eukaryotic gene. Post-processing that considers the nuances of exon, intron, and intergenic length distributions at the gene level could reveal the full potential of this structural approach. Similar information has been part of the success of state-of-the-art tools, both new and old [2–6]. The models trained here would still need to produce exon predictions enriched with true positives for such a post-processing strategy to result in strong performance. That said, it is still likely that genomic features beyond just those around the exon junctions need to be considered for the most robust exon or gene-level predictions. In this vein, another opportunity for improvement could be the inclusion of transcription start sites (TSS) as a class in the deep learning model, as Sharma et al. also found structural signatures at these locations [8]. If separable from others, this class could provide important contextual information for sets of exons, enabling the beginnings of something akin to gene-level predictions. This would require a new training strategy, as TSS profiles were likely lumped into the control class during training. Including TSS profiles in the control set may have negatively affected model performance, especially if they had more in common with boundary profiles than most other exonic and intronic profiles. This is a shortcoming of the labeling strategy used for this work, as broad assumptions had to be made about entire populations of sequences/profiles based on a limited subset of the genomic annotation. There are probably other unutilized subpopulations of structural profiles that are separable from the rest and relevant to this application.

As discussed in **Section 3.4**, ChemEXIN’s performance was even further from acceptable than that of the other 3 models. This was surprising, as the recent paper by Sharma et al. has a table showing all performance metrics having values from ∼0.8 to ∼0.93 in sequences from 3 species, which was much better than those from AUGUSTUS in the same data set [12]. The question naturally arises: what went wrong? As detailed in **Section 2.7**, some of the ChemEXIN source code on GitHub had to be modified for the tool to process more than one sequence at a time. This was necessary to evaluate the tool on the same sequence sets as the other models. After reviewing the source code, it turns out that there are some potential issues with the implementation. Min-max normalization of the profile for an input sequence appears to be conducted based on the minimum and maximum values of the profile, rather than the minimum and maximum values calculated from the training set. While not strictly incorrect, this is not good practice and leaves the model prone to variable performance from one sequence to the next. A viable substitute for this would be to have the model’s input layer include batch normalization, by which it would essentially learn normalization parameters from the training set automatically. Nothing was found in the paper or source code that indicated this was the case for ChemEXIN’s model. There is a batch normalization layer deeper into the network, but it is not positioned such that it could replace the need for typical input normalization (see ***Figure 6*** in the recent paper by Sharma et al.) [12]. The next potential source of ChemEXIN’s poor performance is that the model’s 50 bp input window slides over the profile in 50 bp increments during inference, rather than sliding 1 bp at a time. There are contexts in which this could be acceptable and offer performance benefits. That said, because of how the model was trained, this is analogous to the model only reading 1 out of every 50 pages of a book and trying to summarize the plot. Importantly, this behavior would also put a limit on the number of false positives, meaning the models at the heart of this tool may predict even more false positives than observed in ***Figures 10*** and ***11***. Finally, the model architecture shown in ***Figure 6*** of the paper by Sharma et al., as well as the source code, indicate that a naturally 2D input (50 x 7 matrix) is reshaped into an array of 50 x 7 x 1 x 1 in order for it to be receivable by a 3D convolutional layer [12]. This increases the model complexity without offering a performance benefit and is a choice that implies a fundamental misunderstanding of deep learning architectures. An improved approach would have been to change the convolutional layers to match the shape and meaning of the input data. For example, a 1D convolutional layer would have made more sense for a 50 x 7 input that can be considered a multidimensional sequence or time series.

Some issues discussed above could have impeded the performance of the ChemEXIN command-line tool, even if the models it was built around were robust. This would explain the apparent discrepancy between the evaluation of this tool conducted here and in the paper. Sharma et al. may have focused much more on their model’s performance in the evaluation workflow traditionally used for machine learning than they did on how that success could be translated into a useful bioinformatics tool, if at all. As discussed earlier, how a model’s input is prepared, as well as how its output is processed, can be as important as the model itself. Several of the choices in the source code, model development, and training seem to indicate gaps in understanding about machine learning. For example, the training set used for ChemEXIN, in which exon junctions are highly overrepresented, seems unlikely to prepare any model for reasonable performance in many genomes. A small proportion of the DNA is typically protein-coding, and an even smaller proportion accounts for the exon boundary sites. This motivated a different approach when training the models discussed above. Mishra et al. and Sharma et al. made some interesting and novel observations of features that are probably present in many genomes at an underappreciated level of structural biology: the structure of DNA [7,8,12]. Their proposition that this phenomenon could be useful for *ab initio* detection of genes, and perhaps other features, within newly sequenced genomes is still worth exploring, and deep learning is well-positioned to help achieve this goal. That said, it seems that issues likely arose during this group’s execution of their ideas. The work detailed here offers a glimpse of what an improved implementation might be capable of.

## Code availability

The trained models discussed in this work will be made available on GitHub as part of a command-line tool. For the sake of transparency, the code written to prepare the training set and train these models will also be made available on GitHub “as-is”, with no guarantee of functionality in other environments or systems. Please see the [GitHub repository for this project](https://github.com/GreenMtn-bioinfo/Helix-EXon-finder) for code and updates.

## 5) Supplemental results

### 5.1) Model training times

***Figure 12*** gives a sense of how quickly the 3 model architectures train and make predictions. TCN was the fastest, followed by BiLSTM, and then MBDA-Net. These models were trained on different machines so that training could be conducted in parallel. TCN and MBDA-Net were trained using NVIDIA RTX 3060 (laptop) and RTX 3070 (desktop) GPUs, respectively. BiLSTM was trained on an older system with an NVIDIA GTX 1060. That said, the BiLSTM was RAM-limited during training on the systems with the newer GPUs, so the training time ended up being comparable on the older GPU. As such, ***Figure 12*** still provides a rough impression of the relative speeds of these models. Consistent with statements from Bai et al., the TCN architecture is significantly faster than recurrent ones [21]. The TCN-based model was the largest one, but its training and inference occurred so much more quickly than the other two, which both included recurrent layers, that it still had the lowest total times. As for training efficiency, i.e. time roughly normalized by model size, the ranking was TCN, followed by MBDA-Net, and then BiLSTM. Hardware differences may have affected the ranking order of the latter two models, but as noted above, during trial runs BiLSTM ran at a comparable rate on the new and old GPUs due to access to more RAM on the older system. The results in ***Figure 12*** include the inference times for each epoch, in addition to the time spent actually training. This data could matter when deciding which model to include in a command-line tool. As seen and discussed in the results section, the TCN model had comparable performance to the others. That it makes predictions much more quickly is a valuable and differentiating quality to have. After all, in its intended application, and depending on the sequence length, the model would be expected to make hundreds to millions of predictions in a reasonable amount of time.

**Figure 12:**
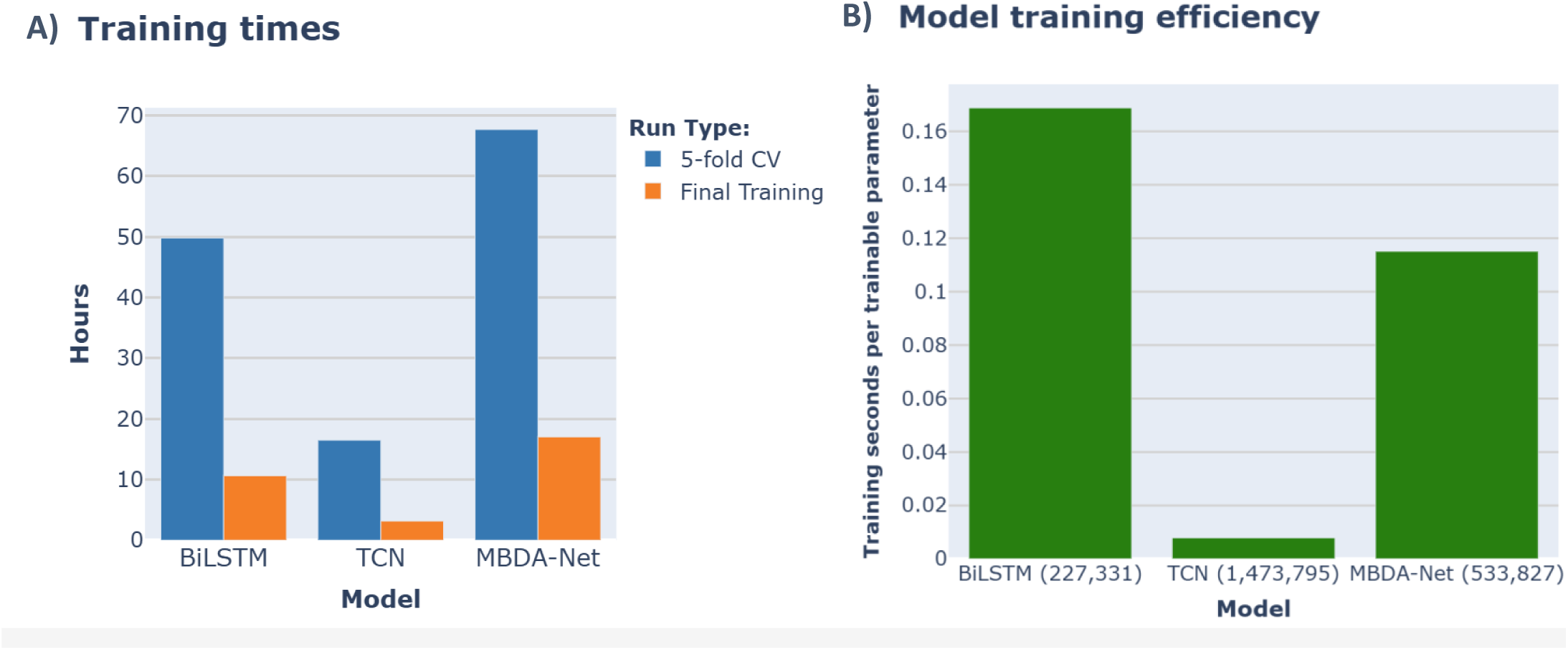
A comparison of the speeds of each model during training and inference. **A)** Total times for 5-fold cross-validation in the training set, as well as the final training (on the full training set) and evaluation (in the test set). **B)** A metric of the efficiency of each model expressed as total seconds spent on the final training divided by its number of trainable parameters (in parentheses next to each model’s name). While the times are referred to as “training times”, they included both the time spent training and inferring/predicting during the epochs.

### 5.2) Cross-validation

***Figure 13*** summarizes the results for all three models from the 5-fold cross-validation in the training set. As noted in **Section 2.4**, cross-validation was run because the hyperparameter tuning process was conducted in a much smaller subset of the training data and was not particularly rigorous or exhaustive. Before proceeding with the evaluation of these models’ performance in the testing set, it seemed worthwhile to ensure the chosen hyperparameters resulted in models that were at least somewhat robust and generalizable. ***Figure 13*** shows that all 3 models reached very similar average performance during cross-validation. The performance achieved later on in the testing set ended up being quite consistent with the average performance seen during cross-validation (compare ***Figures 13*** and ***7***). One difference between the models was their variability when trained on different subsets of the training data and evaluated in others. The BiLSTM model was the most variable, especially in its precision-recall performance (see ***Figure 13A*** and ***13C***). The TCN and MBDA-Net models both varied less, with TCN having the least variable performance. This is consistent with the paper by Bai et al., which explains that the TCN architecture should have more stable gradients than recurrent architectures [21]. Stable gradients mean more predictable/consistent behavior during training, which is more likely to result in models that generalize consistently despite perturbations in the training set. Like its speed, this is another strength of the TCN architecture that may differentiate it for use in a command-line tool, especially in the absence of any significant performance issues compared to the other models.

**Figure 13:**
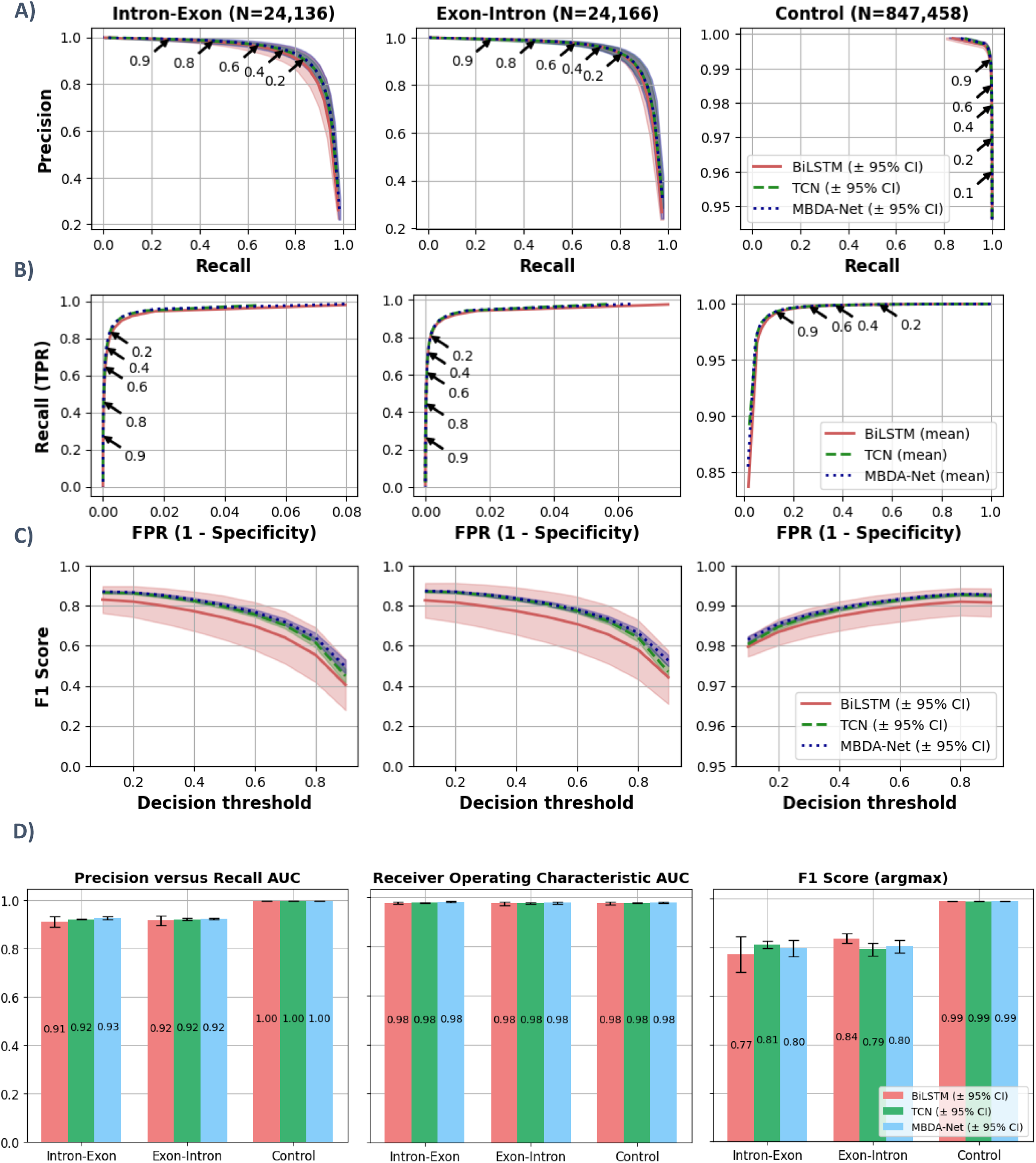
Per-class performance summary from 5-fold cross-validation. Each column of plots in A) - C) corresponds to the class noted at the top, with the number of that class in each validation set in parentheses. Row **A)** contains the precision versus recall (PR) curves. Row **B)** contains the true positive rate (TPR) versus false positive rate (FPR) curves, i.e. the ROC curves. The ±95% confidence interval region was omitted for this row because it was nearly indistinguishable from the mean line. In rows A) and B), decision thresholds are noted for some points on the TCN model’s curve via arrows. Row **C)** contains curves of the F1 Score, i.e. the harmonic mean of precision and recall, at each of the decision thresholds used in the earlier two rows of plots. **D)** Mean area under the curve (AUC) for the precision versus recall curves in A) and the ROC curves in B), as well as the mean F1 score when the class with the max probability is selected (argmax). Some of the plots in a given row are not on the same x and y scales as others.

As discussed in **Section 2.6**, the data expressed in ***Figure 13*** motivated the decision to use a probability cutoff of 0.2 to call either boundary class. While only TCN’s probability thresholds are annotated, for each model a cutoff of ∼0.2 appeared to strike a precision-leaning but still near-optimal balance of precision and recall in the boundary classes (see ***Figure 13A***). The mean false positive rate (FPR) associated with using that threshold was still well below 1%. A caveat is that there are many more controls than boundaries, both in this training set, and in the genome. As such, even relatively small FPRs may translate to large absolute numbers of false positives, which could still make a model unusable. This made the use of an even lower boundary-calling threshold undesirable. As discussed earlier, this model’s output was sent to a post-processing algorithm that performed filtering of predicted boundaries at the exon-level, meaning a decent initial recall at the boundary-level was important. This also influenced the decision to use a cutoff no greater than 0.2. For the control thresholds in ***Figures 13A*** and ***13B***, there is much apparent freedom with regards to achieving strong performance in the control class. Care needed to be taken to regulate the FPR of the control class, and not diminish the recall of the boundary classes too much. The information in ***Figure 13*** could have possibly been used to justify alternative threshold-based filtering schemes at the nucleotide level that resulted in a better-performing post-processing algorithm. That said, the approach outlined in **Section 2.6** was a somewhat effective first attempt. The rightmost plot in ***Figure 13D*** motivated the decision not to use *argmax()* on the model probabilities to call classes, as it resulted in slightly worse F1 scores overall for these models. This was also observed during the development of the post-processing pipeline using longer sequences from the training set. It mostly stemmed from reduced recall in the boundary classes, due to (potential) true positives, with moderate-to-low probability, being classified as a control by *argmax()*.

## References

[1] Wang, Z., Chen, Y., & Li, Y. (2004). A Brief Review of Computational Gene Prediction Methods. Genomics, Proteomics & Bioinformatics, 2(4), 216–221. 10.1016/s1672-0229(04)02028-5.

[2] Stanke, M., Steinkamp, R., Waack, S., & Morgenstern, B. (2004). AUGUSTUS: a web server for gene finding in eukaryotes. Nucleic Acids Research, 32(Web Server), W309–W312. 10.1093/nar/gkh379.

[3] Gabriel, L., Brůna, T., Hoff, K. J., Ebel, M., Lomsadze, A., Borodovsky, M., & Stanke, M. (2024). BRAKER3: Fully automated genome annotation using RNA-seq and protein evidence with GeneMark-ETP, AUGUSTUS, and TSEBRA. Genome Research, 34(5), 769–777. 10.1101/gr.278090.123.

[4] Gabriel, L., Becker, F., Hoff, K. J., & Stanke, M. (2024). Tiberius: end-to-end deep learning with an HMM for gene prediction. Bioinformatics, 40(12). 10.1093/bioinformatics/btae685.

[5] Ye, K., Zhang, P., Xu, T., Wang, S., Yang, X., Sun, P., Jia, P., Wang, B., Bush, S., & Ning, Z. (2025). Highly accurate ab initio gene annotation with ANNEVO. Springer Science and Business Media LLC. 10.21203/rs.3.rs-6402260/v1.

[6] Stiehler, F., Steinborn, M., Scholz, S., Dey, D., Weber, A. P. M., & Denton, A. K. (2020). Helixer: cross-species gene annotation of large eukaryotic genomes using deep learning. Bioinformatics, 36(22–23), 5291–5298. 10.1093/bioinformatics/btaa1044.

[7] Mishra, A., Siwach, P., Misra, P., Dhiman, S., Pandey, A. K., Srivastava, P., & Jayaram, B. (2021). Intron exon boundary junctions in human genome have in-built unique structural and energetic signals. Nucleic Acids Research, 49(5), 2674–2683. 10.1093/nar/gkab098.

[8] Sharma, D., Sharma, K., Mishra, A., Siwach, P., Mittal, A., & Jayaram, B. (2023). Molecular dynamics simulation-based trinucleotide and tetranucleotide level structural and energy characterization of the functional units of genomic DNA. Physical Chemistry Chemical Physics, 25(10), 7323–7337. 10.1039/d2cp04820e.

[9] Nieto Moreno, N., Giono, L. E., Cambindo Botto, A. E., Muñoz, M. J., & Kornblihtt, A. R. (2015). Chromatin, DNA structure and alternative splicing. In FEBS Letters (Vol. 589, Issue 22, pp. 3370–3378). Wiley. 10.1016/j.febslet.2015.08.002.

[10] Tsai, Z. T.-Y., Chu, W.-Y., Cheng, J.-H., & Tsai, H.-K. (2013). Associations between intronic non-B DNA structures and exon skipping. In Nucleic Acids Research (Vol. 42, Issue 2, pp. 739–747). Oxford University Press (OUP). 10.1093/nar/gkt939.

[11] Wang, G., & Vasquez, K. M. (2022). Dynamic alternative DNA structures in biology and disease. In Nature Reviews Genetics (Vol. 24, Issue 4, pp. 211–234). Springer Science and Business Media LLC. 10.1038/s41576-022-00539-9.

[12] Sharma, D., Aslam, D., Sharma, K., Mittal, A., & Jayaram, B. (2025). Exon–intron boundary detection made easy by physicochemical properties of DNA. Molecular Omics, 21(3), 226–239. 10.1039/d4mo00241e.

[13] Kopka, M. L., Fratini, A. V., & Dickerson, R. E. (1982). REVERSIBLE BENDING AND HELIX GEOMETRY IN A B-DNA DODECAMER: CGCGAATTBRCGCG [Dataset]. In Worldwide Protein Data Bank. Worldwide Protein Data Bank. 10.2210/pdb4bna/pdb.

[14] Lu, X.-J., Bussemaker, H. J., & Olson, W. K. (2015). DSSR: an integrated software tool for dissecting the spatial structure of RNA. Nucleic Acids Research, gkv716. 10.1093/nar/gkv716.

[15] Da Rosa, G., Grille, L., Calzada, V., Ahmad, K., Arcon, J. P., Battistini, F., Bayarri, G., Bishop, T., Carloni, P., Cheatham III, T., Collepardo-Guevara, R., Czub, J., Espinosa, J. R., Galindo-Murillo, R., Harris, S. A., Hospital, A., et al. (2021). Sequence-dependent structural properties of B-DNA: what have we learned in 40 years? In Biophysical Reviews (Vol. 13, Issue 6, pp. 995–1005). Springer Science and Business Media LLC. 10.1007/s12551-021-00893-8.

[16] Genome Reference Consortium. (2022). Human build 38 patch release 14 (GRCh38.p14) (Accession Nos. GCA_000001405.29, GCF_000001405.40) [Dataset]. National Center for Biotechnology Information. https://www.ncbi.nlm.nih.gov/assembly/11968211.

[17] Goldfarb, T., Kodali, V. K., Pujar, S., Brover, V., Robbertse, B., Farrell, C. M., Oh, D.-H., Astashyn, A., Ermolaeva, O., Haddad, D., et al. (2024). NCBI RefSeq: reference sequence standards through 25 years of curation and annotation. Nucleic Acids Research, 53(D1), D243–D257. 10.1093/nar/gkae1038.

[18] Mokry, M., Feitsma, H., Nijman, I. J., de Bruijn, E., van der Zaag, P. J., Guryev, V., & Cuppen, E. (2010). Accurate SNP and mutation detection by targeted custom microarray-based genomic enrichment of short-fragment sequencing libraries. Nucleic Acids Research, 38(10), e116–e116. 10.1093/nar/gkq072.

[19] Danecek, P., Bonfield, J. K., Liddle, J., Marshall, J., Ohan, V., Pollard, M. O., Whitwham, A., Keane, T., McCarthy, S. A., Davies, R. M., & Li, H. (2021). Twelve years of SAMtools and BCFtools. GigaScience, 10(2). 10.1093/gigascience/giab008.

[20] Rémy, P. (2021). keras-tcn [Computer software]. GitHub. https://github.com/philipperemy/keras-tcn.

[21] Bai, S., Kolter, J. Z., & Koltun, V. (2018). An Empirical Evaluation of Generic Convolutional and Recurrent Networks for Sequence Modeling (Version 2). arXiv. 10.48550/ARXIV.1803.01271.

[22] Gao, J., Gao, X., Wu, N., & Yang, H. (2022). Bi-directional LSTM with multi-scale dense attention mechanism for hyperspectral image classification. Multimedia Tools and Applications, 81(17), 24003–24020. 10.1007/s11042-022-12809-z.

[23] Amidi, A., & Amidi, S. (2018). A detailed example of how to use data generators with Keras. Afshine Amidi’s Blog. Standford University. https://stanford.edu/∼shervine/blog/keras-how-to-generate-data-on-the-fly.

[24] xor2k, synapticarbors [Adelman, J.], & gityoav [Git, Y.] (2025). npy-append-array [Computer software]. GitHub. https://github.com/xor2k/npy-append-array.

[25] Aslam, D., Sharma, D., & Sharma, K. (2024). ChemEXIN [Computer software]. GitHub. https://github.com/rnsharma478/ChemEXIN.

[26] Movassat, M., Forouzmand, E., Reese, F., & Hertel, K. J. (2019). Exon size and sequence conservation improves identification of splice-altering nucleotides. RNA, 25(12), 1793–1805. 10.1261/rna.070987.119.

